# Dietary intake regulates the circulating inflammatory monocyte pool

**DOI:** 10.1101/582346

**Authors:** Stefan Jordan, Navpreet Tung, Maria Casanova-Acebes, Christie Chang, Claudia Cantoni, Dachuan Zhang, Theresa H. Wirtz, Shruti Naik, Samuel A. Rose, Chad N. Brocker, Anastasiia Gainullina, Barbara B. Maier, Derek LeRoith, Frank J. Gonzalez, Felix Meissner, Jordi Ochando, Adeeb Rahman, Jerry E. Chipuk, Maxim N. Artyomov, Paul S. Frenette, Laura Piccio, Marie-Luise Berres, Emily J. Gallagher, Miriam Merad

**Affiliations:** Department of Oncological Sciences, Icahn School of Medicine at Mount Sinai, 1 Gustave L. Levy Place, New York, NY 10029, USA; The Precision Immunology Institute, Icahn School of Medicine at Mount Sinai, 1 Gustave L. Levy Place, New York, NY 10029, USA; The Tisch Cancer Institute, Icahn School of Medicine at Mount Sinai, 1 Gustave L. Levy Place, New York, NY 10029, USA; Department of Genetics and Genomic Sciences, Icahn School of Medicine at Mount Sinai, 1 Gustave L. Levy Place, New York, NY 10029, USA; Division of Endocrinology, Diabetes and Bone Diseases, Icahn School of Medicine at Mount Sinai, 1 Gustave L. Levy Place, New York, NY 10029, USA; Human Immune Monitoring Center, Icahn School of Medicine at Mount Sinai, 1 Gustave L. Levy Place, New York, NY 10029, USA; Ruth L. and David S. Gottesman Institute for Stem Cell and Regenerative Medicine Research, Department of Cell Biology, Albert Einstein College of Medicine, 1301 Morris Park Avenue, The Bronx, NY 10461, USA.; Department of Internal Medicine III, University Hospital, RWTH Aachen, Pauweisstrasse 30, 52074 Aachen, Germany; Department of Pathology, and Ronald O. Perelman Department of Dermatology, NYU School of Medicine, 240 East 38^th^ Street, New York, NY 10016, USA; Laboratory of Metabolism, Center for Cancer Research, National Cancer Institute, National Institutes of Health, Building 37, Bethesda, MD 20892, USA; Department of Pathology & Immunology, Washington University School of Medicine, 660 S Euclid Avenue, St. Louis, MO 63110, USA; Department of Neurology, Washington University School of Medicine, 660 S Euclid Avenue, St. Louis, MO 63110, USA; Computer Technologies Department, ITMO University, Kronverksky 49, Saint Petersburg, Russian Federation; Max-Planck-Institute of Biochemistry, Am Klopferspitz 18, 82152 Martinsried, Germany

**Keywords:** Caloric restriction, fasting, metabolism, inflammation, monocyte, liver, AMPK, PPARα, CCL2, inflammatory disease

## Abstract

Caloric restriction is known to improve inflammatory and autoimmune diseases. However, the mechanisms by which reduced caloric intake modulates inflammation are poorly understood. Here we show that short-term fasting reduced monocyte metabolic and inflammatory activity and drastically reduced the number of circulating monocytes. Regulation of peripheral monocyte numbers was dependent on dietary glucose and protein levels. Specifically, we found that activation of the low-energy sensor 5’-AMP-activated protein kinase (AMPK) in hepatocytes and suppression of systemic CCL2 production by peroxisome proliferator-activator receptor alpha (PPARα) reduced monocyte mobilization from the bone marrow. Importantly, while caloric restriction improves chronic inflammatory diseases, fasting did not compromise monocyte emergency mobilization during acute infectious inflammation and tissue repair. These results reveal that caloric intake and liver energy sensors dictate the blood and tissue immune tone and link dietary habits to inflammatory disease outcome.

**Highlights:**

- Fasting reduces the numbers of peripheral pro-inflammatory monocytes in healthy humans and mice.
- A hepatic AMPK-PPARα energy-sensing axis controls homeostatic monocyte numbers via regulation of steady-state CCL2 production.
- Fasting reduces monocyte metabolic and inflammatory activity.
- Fasting improves chronic inflammatory diseases but does not compromise monocyte emergency mobilization during acute infectious inflammation and tissue repair.

## INTRODUCTION

Caloric excess, which is frequent in diets in the western world, has been linked to systemic low-grade chronic inflammation (Lumeng and Saltiel, 2011), and is thought to contribute to numerous diseases including metabolic syndrome (MetS), non-alcoholic fatty liver disease (NAFLD), type 2 diabetes mellitus (T2DM), atherosclerosis, cardiovascular disease (CVD), and other related co-morbidities (Haslam and James, 2005). Accordingly, the recent diet westernization of developing countries has been associated with an increased prevalence of inflammatory or autoimmune disorders (Manzel et al., 2014). In contrast, reduced caloric intake is associated with improved outcomes for metabolic, autoimmune and inflammatory diseases, including NAFLD (Kani et al., 2017), T2DM (Cheng et al., 2017), CVD (Wei et al., 2017), multiple sclerosis (Choi et al., 2016; Jahromi et al., 2014), rheumatoid arthritis (Choi et al., 2017; Kjeldsen-Kragh et al., 1991), asthma (Johnson et al., 2007), psoriasis (Jensen et al., 2016), colitis (Shibolet et al., 2002), and sepsis (Starr et al., 2016; Wannemacher et al., 1979) and has been shown to prolong the lifespan of rodents, monkeys, and humans (Fontana et al., 2010; Picca et al., 2017). However, the molecular mechanisms by which caloric restriction modulates systemic inflammation remain poorly understood.

Clinical studies performed in overweight or obese individuals undergoing caloric restriction showed a reduction of pro-inflammatory cytokines in the blood (Ho et al., 2015; Ikizler et al., 2018; Imayama et al., 2012; Loria-Kohen et al., 2013; Oh et al., 2013; Ott et al., 2017; Ramel et al., 2010; Tajik et al., 2013), and diet-induced weight loss has a superior benefit on patient systemic inflammation compared to interventional weight loss due to gastric bypass surgery (Lips et al., 2016). Although little data is available on the effect of caloric restriction on inflammation in normal-weight individuals, prior studies have shown that individuals undergoing intermittent or religious fasting have reduced basal levels of circulating pro-inflammatory cytokines including TNFα, IL-6, and IL-1β (Aksungar et al., 2007; Faris et al., 2012; Moro et al., 2016).

Prompted by these prior results, we sought to investigate the impact of fasting on immune cell homeostasis. We first used mass cytometry to profile blood circulating cells of healthy, normal-weight humans prior to and during fasting. Strikingly, we discovered that fasting significantly reduced the number of circulating monocytes and similar results were obtained in mice. Here we describe how dietary energy intake controls the quality and quantity of blood and tissue monocytes emphasizing the link between high calorie dietary patterns and inflammatory disease outcome in patients.

## RESULTS

### Fasting Reduces the Pool of Circulating Monocytes in Healthy Humans and Mice

To explore whether fasting was associated with changes in peripheral blood leukocyte populations, we profiled the composition of blood circulating immune cells of 12 healthy normal weight volunteers (mean age = 30+/-5 years, BMI = 22+/-2 kg/m^2^) 3 hrs after food intake (fed state) and after 19 hrs of fasting (fasting state) using Cytometry by Time-Of-Flight spectrometry (CyTOF) (Figures 1A and 1B). To control for circadian variations, all blood samples were drawn at the same time of the day (3 pm). Strikingly, we found that fasting led to significant reduction of circulating monocytes including both CD14+ and CD16+ monocyte subsets (Figures 1C, 1D and S1A). Interestingly, in individuals with low baseline monocyte numbers, fasting did not decrease monocyte numbers below the physiologic range (Figure 1D). In addition to monocytes, a small subset of circulating dendritic cells, called CD141+ DC, was also reduced, whereas neutrophils were not significantly affected during short term fasting (Figure 1E).

**Figure 1.**
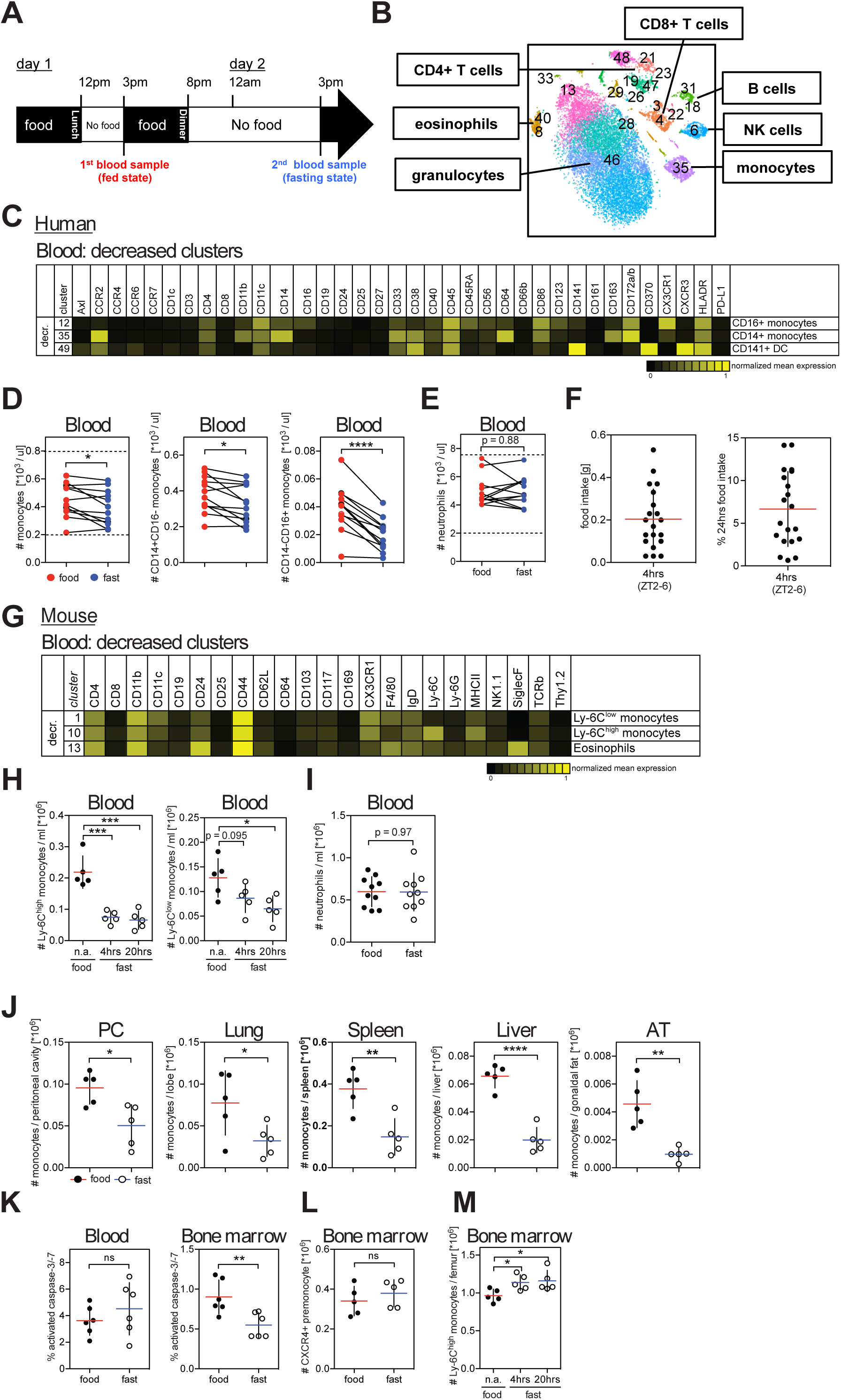
Fasting Reduces the Number of Circulating Pro-inflammatory Monocytes in Healthy Humans and Mice. (A) Schematic representation of the fasting experimental design. (B to E) Blood was drawn from healthy individuals in the fed and in the fasting state and analyzed by CyTOF. (B) Multidimensional CyTOF data were clustered using viSNE. (C) Heatmap shows mean markers expression on cell clusters significantly reduced during fasting. (D and E) Paired analysis of (D) total monocytes, CD14+CD16-monocytes, and CD14-CD16+ monocytes, and (E) neutrophils in human blood during the fed and the fasted state. Dotted lines indicate physiologic range. (F) Food intake of individual mice during 4 hrs between ZT2 and ZT6, and percentage of food intake between ZT2 and ZT6 with regard to 24 hr food intake. (G) CyTOF analysis of blood cells from fed and short-term fasted mice. Heatmap shows mean marker expression of clusters significantly reduced by fasting. (H) Absolute numbers of Ly-6C^high^ and Ly-6C^low^ monocytes in the blood of mice that were fed or fasted for the indicated time. (I) Absolute numbers of neutrophils in the blood of mice that were fed or fasted for 20 hrs. (J) Absolute numbers of Ly-6C^high^ monocytes in the peritoneal cavity (PC), lung, spleen, liver and adipose tissue (AT) of mice that were fed or fasted for 20 hrs. (K) Percentage of caspase-3/7+ cells among Ly-6C^high^ monocytes in the blood and bone marrow of mice that were fed or fasted for 4 hrs. (L) Numbers of bone marrow CXCR4+ pre-monocytes in mice that were fed or fasted for 20 hrs. (M) Absolute numbers of bone marrow Ly-6C^high^ monocytes in mice that were fed or fasted for the indicated time. (F, H to M) Every dot represents one individual animal. Horizontal bar = mean. Vertical bar = SD. Student’s t test (D, E, I to L) or one-way analysis of variance (ANOVA) with Dunnett’s multiple comparison test (H, M) were performed. Statistical significance is indicated by *P < 0.05, **P < 0.01, ***P < 0.001, ****P < 0.0001. ns = not significant. See also Figure S1.

We then asked whether similar changes in blood monocyte numbers also occurred in fasting mice. We chose a 4 hrs short-term fasting protocol during the light period (Zeitgeber [ZT]2-6) which is comparable to overnight fasting for humans and is the least stressful fasting strategy in animals (Figure S1B) (Jensen et al., 2013). Within 4 hrs, mice eat up to 0.5g of chow (0.2g on average) representing ∼7% of their total 24 hrs food intake (Figure 1F). Intriguingly, while the frequencies of most immune cell clusters were unaffected by short-term fasting, Ly-6C^high^ pro-inflammatory monocytes were strongly reduced (Figures 1G and 1H). Ly-6C^low^ patrolling monocytes were proportionally reduced at 4 hrs and reduction in absolute numbers reached significance after 20 hrs of fasting. Prolonged fasting periods (ZT10-6) also reduced additional peripheral leukocyte populations including eosinophils, NK cells and T cells (Figure S1C). Importantly, fasting also led to significant reduction of pro-inflammatory Ly-6C^high^ monocytes in peripheral tissues including the peritoneal cavity, lung, liver, spleen and adipose tissues (Figures 1J, S1D and S1E).

Decreased numbers of circulating monocytes in fasting mice could be due to increased monocyte cell death, reduced bone marrow (BM) myelopoiesis, or reduced BM egress to the periphery. We did not detect an increased number of activated caspase-positive monocytes suggesting that the reduction of peripheral monocytes in fasting mice may not be due to increased cell death (Figure 1K). We also failed to detect decreased numbers of BM CXCR4+ monocyte precursors (Chong et al., 2016) (Figure 1L). Instead, Ly-6C^high^ monocytes accumulated in the BM of fasting mice (Figure 1M). These results suggest that fasting– induced reduction of blood monocytes is due to reduced monocyte egress from the BM to the blood circulation. An important mechanism controlling BM cell egress is neuronal stimulation of β3 adrenergic receptors (β3-AR) on BM stromal cells, which leads to reduced BM CXCL12 production and release of hematopoietic cells into the blood circulation (Mendez-Ferrer et al., 2008). Injection of the β3-AR agonist CL 316,243 into fasting mice partially rescued monocyte egress, but never to the same extent as in fed mice, indicating the existence of additional mechanisms suppressing egress during fasting (Figure S1F). CXCL12 BM levels were not affected by fasting suggesting that CXCL12 does not mediate fasting-induced monocyte accumulation in the BM (Figure S1G). In addition, circadian fluctuations in monocyte release or forced shifts in circadian rhythm (jetlag) did not affect fasting ability to modulate BM monocyte egress (Figures S1H and S1I). Re-feeding mice for 4hrs after an overnight fast restored monocyte numbers in the periphery (Figure S1J), showing that fasting-induced inhibition of BM egress is revoked upon food intake.

### AMPK-mediated Sensing of Dietary Energy Levels Controls the Size of the Peripheral Monocyte Pool

Fasting deprives humans and experimental animals of macronutrients (digestible carbohydrates, protein, fat) as well as essential micronutrients (vitamins, minerals, non-digestible fiber). Removal of macronutrients from the diet was sufficient to reduce the pool of circulating monocytes (Figure 2A). Conversely, oral gavage of fasting mice with isocaloric amounts of carbohydrates and proteins, but not with fat, rescued monocyte numbers in the blood (Figure 2B). Importantly, the size of the monocyte pool in the blood circulation depended on the amount of carbohydrate ingested (Figure 2C).

**Figure 2.**
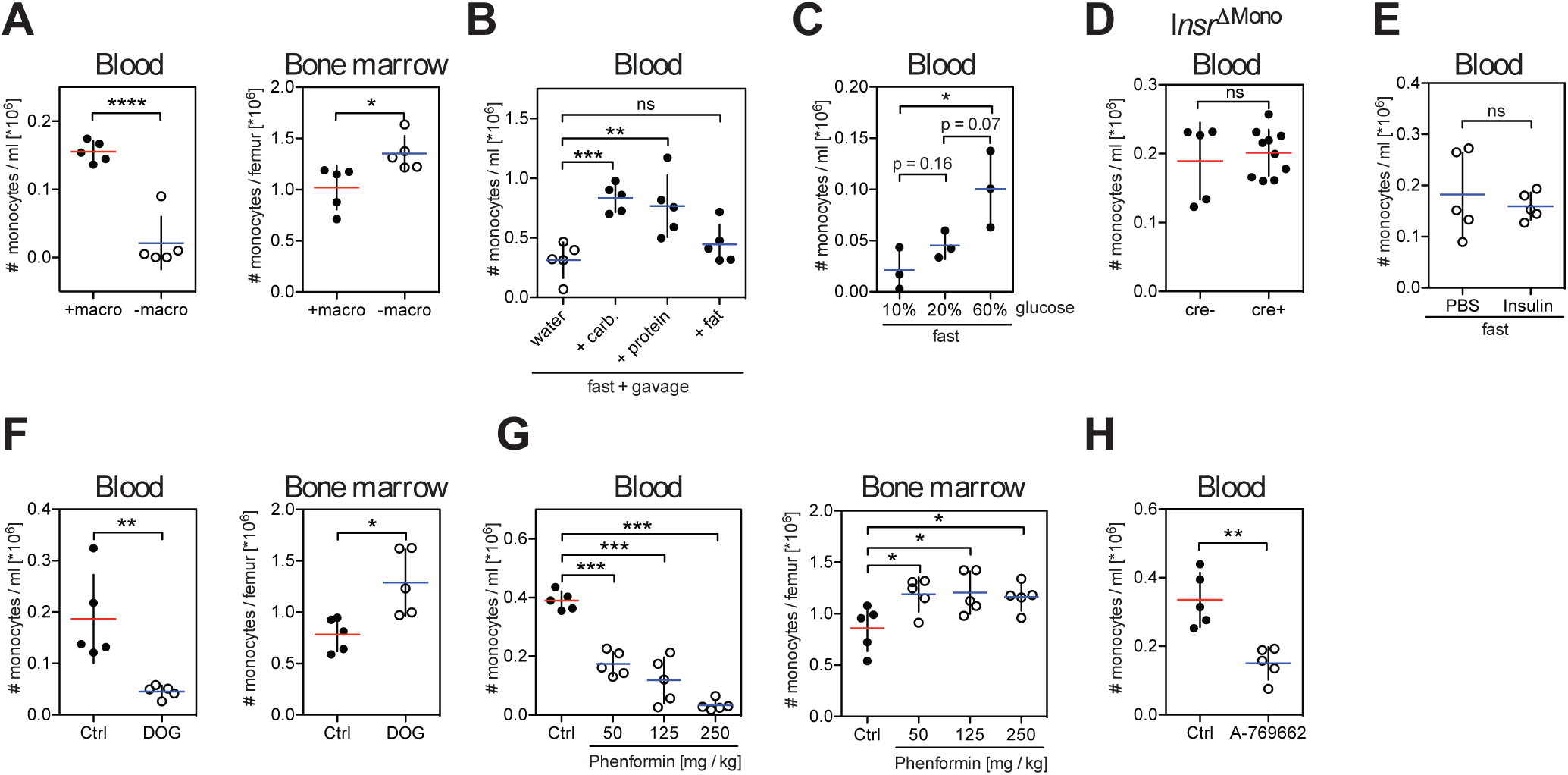
The Energy-sensor AMP-activated Protein Kinase (AMPK) Controls Blood Monocyte Homeostasis. (A to H) Absolute numbers of Ly-6C^high^ monocytes (A) in the blood and bone marrow of mice fed with a diet with (+macro) or without (-macro) macronutrients, (B) in the blood of fasting mice gavaged with water, isocaloric amounts of carbohydrates, protein or fat for 4 hrs, (C) in the blood of mice fasted for 16 hrs and gavaged with glucose solutions at the indicated concentrations, (D) in the blood of fed mice in which the insulin receptor has been deleted from monocytes 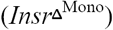, (E) in the blood of mice that were fasted for 4 hrs and injected with insulin 30 minutes prior to analysis, (F) in the blood and bone marrow of mice that were gavaged with water (Ctrl) or 2-deoxyglucose (DOG) once every hour for 4 hrs, (G) in the blood and bone marrow of mice that received water (Ctrl) or a single dose of phenformin 4 hrs before assessment, (H) in blood of mice that were gavaged with AMPK activator A-769662 4 hrs before analysis. (A to H) Every dot represents one individual animal. Horizontal bar = mean. Vertical bar = SD. One-way analysis of variance (ANOVA) with Tukey’s multiple comparison test (C) or Dunnett’s test (B, G), or Student’s t test (A, D to F, H) were performed. Statistical significance is indicated by *P < 0.05, **P < 0.01, ***P < 0.001, ****P < 0.0001. ns = not significant. See also Figure S2.

Carbohydrate and protein intake stimulate insulin secretion, prompting us to examine the contribution of insulin in the regulation of BM monocyte egress. However, we found that genetic deletion of the insulin receptor gene in monocytes (Figure 2D) or exogenous insulin administration in fasting mice (Figure 2E) did not affect peripheral blood monocyte numbers suggesting that insulin was not responsible for BM monocyte egress. Therefore, we hypothesized that carbohydrates and proteins might modulate peripheral monocyte numbers by altering cellular energy levels. To address this hypothesis, we used two different inhibitors of hexokinase, 2-deoxyglucose and D-mannoheptulose, in order to block the first step in glycolysis, i.e. cellular energy production (Figures 2F and S2). Interestingly, blocking glycolysis reduced monocyte numbers to levels similar to those observed during fasting, suggesting that cellular energy levels control the blood monocyte pool.

Mammalian 5’-AMP-activated protein kinase (AMPK) is a key cellular energy sensor triggered by an increase in the cellular AMP/ATP ratio that reflects low energy levels. Phenformin elevates the cellular AMP/ATP ratio resulting in AMPK activation. Strikingly, we found that phenformin administration significantly reduced monocyte egress in a dose-dependent manner (Figure 2G). To further examine the contribution of AMPK to the regulation of blood monocyte levels we gavaged mice with a small molecule activator of AMPK (A-769662) (Figure 2H). Consistent with the data obtained with phenformin administration, we found that oral gavage with A-769662 significantly reduced the pool of blood peripheral monocytes in fed mice. Altogether these results suggest that activation of the low-energy sensor AMPK is sufficient to inhibit BM monocyte egress to the blood circulation.

### Hepatic PPARα Controls Peripheral Monocyte Numbers

Peroxisome proliferator-activated receptor α (PPARα) is a target of several nutrient-sensing pathways including AMPK and is a master transcriptional regulator in the adaptive response to fasting. PPARα has been implicated in the anti-inflammatory effects of fasting (Wang et al., 2016; Youm et al., 2015). Strikingly, we found that fasting-induced reduction of circulating monocytes was less efficient in *Ppara*^−/−^ mice compared to wild-type mice suggesting that activation of PPARα contributed to the regulation of monocyte homeostasis during fasting (Figure 3A). Phenformin treatment had a much more pronounced effect in *Ppara*^+/+^ mice compared to *Ppara*^−/−^ mice, indicating that AMPK-mediated reduction in peripheral monocyte numbers in fasting mice was in part mediated through PPARα (Figure 3B).

**Figure 3.**
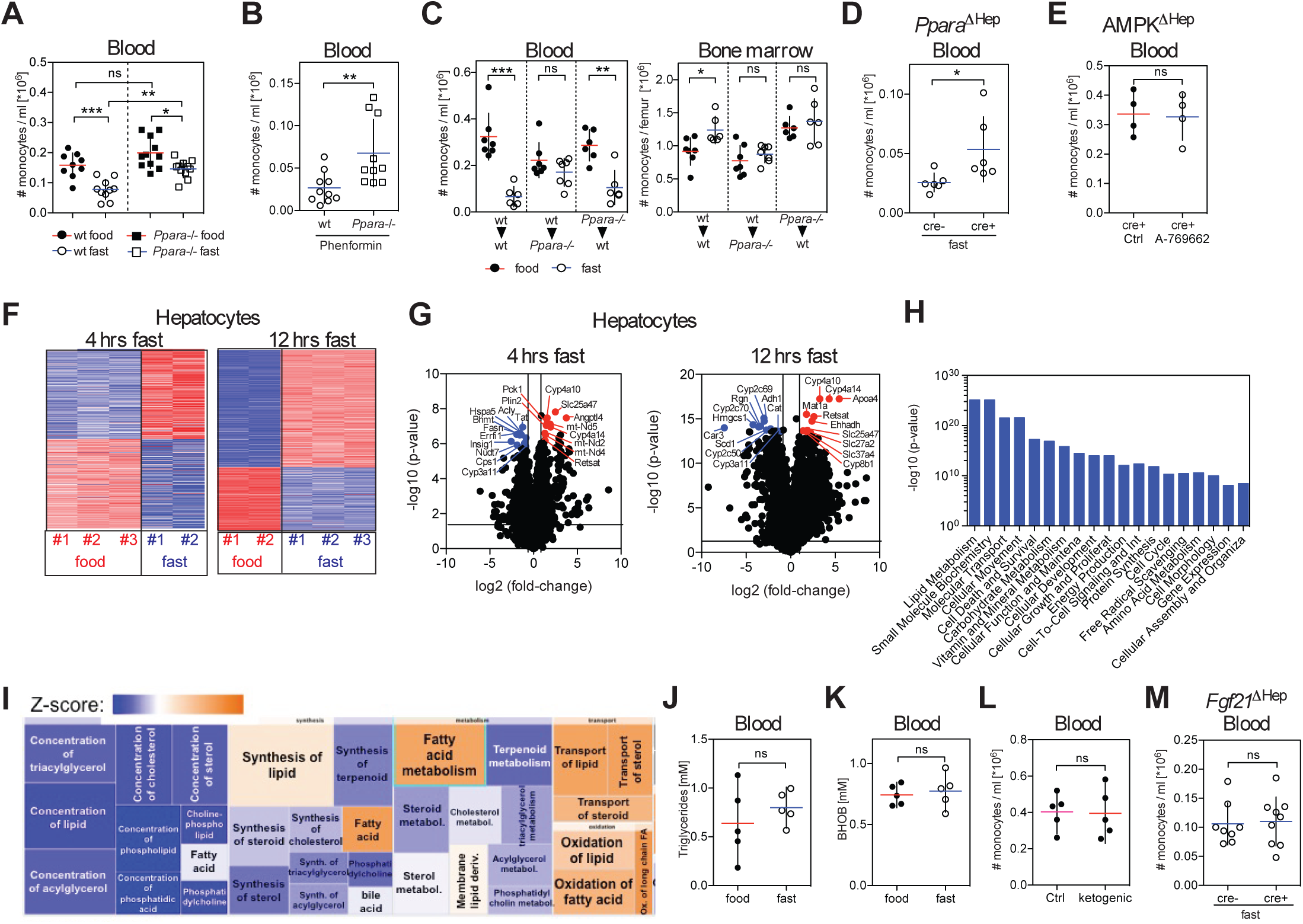
Liver Fasting Metabolism Regulator PPARα Controls Peripheral Monocyte Numbers in the Steady State. (A and B) Absolute numbers of Ly-6C^high^ monocytes were measured (A) in the blood of wild-type and *Ppara*^−/−^ mice that were fed or fasted for 4 hrs, and (B) in the blood of wild-type and *Ppara*^−/−^ mice gavaged with phenformin 4 hrs prior to analysis. (C) Bone marrow chimeric mice were generated so that wild-type mice (wt) were reconstituted with wt or *Ppara*^−/−^ bone marrow cells, and *Ppara*^−/−^ mice were reconstituted with wt bone marrow cells. Seven weeks after reconstitution half of the mice from each group were fasted and the absolute numbers of Ly-6C^high^ monocytes in the blood and bone marrow of each bone marrow chimeric group were measured. (D) *Alb*^cre/cre^ mice were crossed to *Ppara*^fl/fl^ mice to delete PPARα from hepatocytes (*Ppara***Δ**^Hep^) in cre+ mice. Graph shows absolute numbers of Ly-6C^high^ monocytes in the blood circulation of cre- and cre+ mice after 20 hrs of fasting. (E) *Alb*^cre/cre^ mice were crossed to *Prkaa1*^fl/fl^ mice to delete AMPK from hepatocytes (*AMPK***Δ**^Hep^). Numbers of Ly-6C^high^ monocytes in blood of cre+ mice that were gavaged with control solvent or A-769662 4 hrs before analysis are shown. (F) Heatmap shows z-scores of differentially expressed genes in hepatocytes after 4 hrs and 12 hrs of fasting. (G) Volcano plot identifies 10 most up- and downregulated transcripts in hepatocytes at 4 hrs and 12 hrs of fasting. (H and I) Ingenuity pathway analysis (IPA) of differential expressed genes in hepatocytes after 4 hrs of fasting. (H) Bar chart shows most significantly altered cellular functions. (I) Detailed view of molecular functions within lipid metabolism that are affected during fasting. (J and K) Levels of (J) triglycerides and (K) β-hydroxybutyrate (BHOB) in the blood of mice that were fed or fasted for 4 hrs. (L) Ly-6C^high^ monocytes in blood of mice that were fed with ketogenic or control diet for 3 weeks. (M) *Alb*^cre/cre^ mice were crossed to *Fgf21*^fl/fl^ mice to delete *Fgf21* from hepatocytes 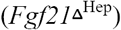 and cre-or cre+ mice were fasted for 4 hrs. Graph shows absolute numbers of Ly-6C^high^ monocytes in the blood. (A) Data were pooled from two experiments. (A to E, J to M) Every dot represents one individual animal. Horizontal bar = mean. Vertical bar = SD. One-way analysis of variance (ANOVA) with Bonferroni’s test (A), or Student’s t test (B to E, J to M) were performed. Statistical significance is indicated by *P < 0.05, **P < 0.01, ***P < 0.001. ns = not significant. See also Figure S3.

To determine whether PPARα mediated its effect on immune cell homeostasis by acting directly in monocytes, we generated bone marrow chimeric animals in which wild-type or *Ppara*^−/−^ bone marrow cells were injected into lethally irradiated hosts (Figure 3C). Interestingly, whereas monocyte egress was reduced in fasting wild-type mice reconstituted with *Ppara*^−/−^ BM, we failed to observe a significant reduction of blood monocytes in fasting *Ppara*^−/−^ mice reconstituted with wild-type BM, thus establishing that the control of BM monocyte egress required PPARα expression in non-hematopoietic cells and not in monocytes.

PPARα is expressed at higher levels in the liver (Figure S3A) and acts mainly in hepatocytes (Brocker et al., 2017), prompting us to probe the role of hepatic PPARα in the control of peripheral monocyte numbers. Strikingly, we found that mice in which PPARα was deleted uniquely in hepatocytes were impaired in their ability to modulate BM monocyte egress upon fasting (Figure 3D). Since modulation of BM monocyte egress upon fasting required hepatic PPARα, we tested whether sensing of low-energy levels by AMPK also occurred in hepatocytes. Importantly, we found that deletion of AMPK specifically in hepatocytes abrogated the reduction of circulating monocyte numbers upon gavage with an AMPK activator (Figure 3E). Altogether, these data establish that energy-sensing by the liver AMPK-PPARα pathway controls the blood monocyte pool in response to caloric intake. Consistent with this idea, transcriptional analysis of hepatocytes isolated after 4 hrs and 12 hrs of fasting revealed that most differentially expressed genes were involved in metabolic processes (Figures 3F and 3G) with the top transcriptional regulator at both time-points being PPARα (p-value 3.09×10^−25^ and 1.16×10^−46^ at 4 hrs and 12 hrs, respectively). Genes involved in lipid metabolism were strongly modulated upon short-term fasting (p-value range from 4×10^−6^-1.27×10^−25^) (Figures 3H and 3I) prompting us to examine whether metabolic adaptation to the fasting state played a role in the suppression of monocyte egress. However, we found that global triglyceride and ketone body levels as well as levels of most individual metabolites tested were unaffected by short-term fasting (Figures 3J, 3K, S3B and S3C). In addition, a ketogenic diet driving lipid metabolism failed to affect peripheral monocyte numbers (Figure 3L). Finally, we interfered with fasting-induced lipid mobilization from fat depots by deleting FGF21 expression in liver, but could not detect any changes in monocyte numbers in fasting *Fgf21*^−/−^ mice compared to littermate controls (Figure 3M). In summary, these data suggested that lipid metabolites do not play a major role in the regulation of BM monocyte egress.

### PPARα Controls Steady-State CCL2 Levels

Transition from the fed to the fasting state involves changes in the plasma levels of numerous metabolic hormones. Short-term fasting induced remarkably similar changes in the plasma hormonal profile of humans and mice with increased levels of ghrelin and decreased levels of insulin, c-peptide, amylin, GIP, leptin, PP, and PYY (Figure 4A and S4A). Most interestingly, CCL2 (also known as MCP-1) plasma level was reduced in 8 out of 12 fasting human subjects. CCL2 binds to CCR2, a chemotactic receptor highly expressed on monocytes and shown to mediate monocyte BM egress (Serbina and Pamer, 2006). We also found a strong reduction of systemic CCL2 in fasting mice (Figures 4A and 4B) as well as upon administration of AMPK-activator phenformin (Figure 4C). Importantly, restoring plasma CCL2 levels by administration of recombinant protein rescued monocyte numbers in fasting mice (Figure 4D), further indicating the critical role for CCL2 in monocyte homeostasis during fasting. Providing glucose in fasting *Ccr2*-/- mice failed to elevate blood monocyte numbers and thus did not bypass *Ccr2*-deficiency (Figure S4B).

**Figure 4.**
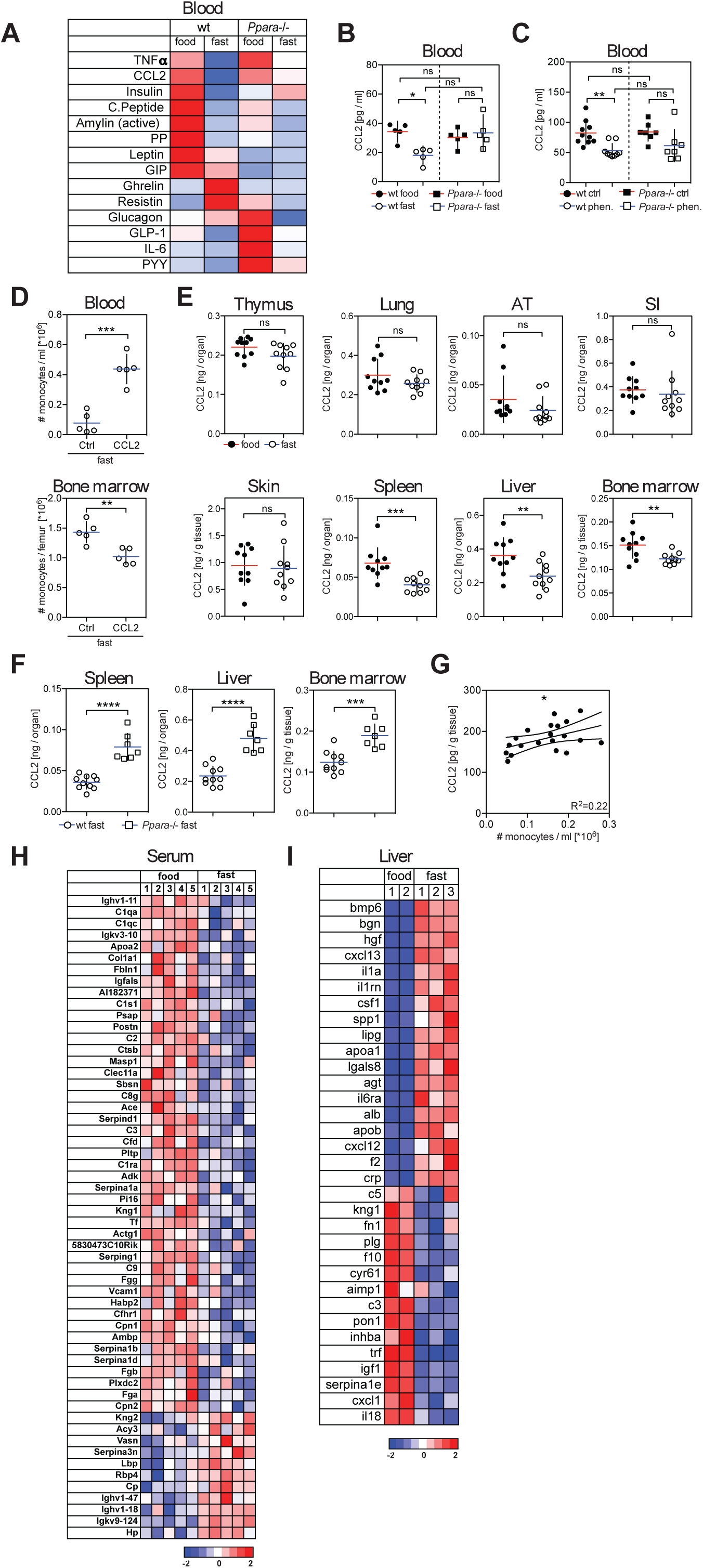
PPARα Controls Tissue CCL2 Levels in the Steady-State. (A) Multiplex analysis for metabolic hormones in mouse blood in the fed and fasting state. (B) CCL2 levels in blood of wild-type (wt) and *Ppara*^−/−^ mice that were fed or fasted for 4 hrs. (C) CCL2 levels in blood of wild-type (wt) and *Ppara*^−/−^ mice that were gavaged with water (Ctrl) or with a single dose of phenformin 4 hrs before analysis. Data were pooled from two experiments. (D) Mice were fasted for 20 hrs before injection of PBS or recombinant CCL2 and were analyzed 3 hrs after injection. Graphs show the absolute numbers of Ly-6C^high^ monocytes in the blood and bone marrow. (E) CCL2 protein in indicated tissues in fed and fasted wt mice. AT = adipose tissue, SI = small intestine. (F) CCL2 protein in the indicated tissues in fasted wt and *Ppara*^−/−^ mice. (G) Plot shows CCL2 production in BM vs. monocyte numbers in blood. (H) Heatmap shows z-scores of significantly changing serum protein levels in fed and fasted mice. (I) Differentially expressed genes in liver upon fasting were filtered using the IPA database. Heatmap shows z-scores of genes that are known to regulate CCL2 production and gene products are secreted to the extracellular space. (B to G) Every dot represents one individual animal. Horizontal bar = mean. Vertical bar = SD. One-way analysis of variance (ANOVA) with Tukey’s post test (B) or Bonferroni’s post test (C), or Student’s t test (D to F) were performed. Statistical significance is indicated by *P < 0.05, **P < 0.01, ***P < 0.001, ****P < 0.0001. ns = not significant. See also Figure S4.

CCL2 is produced in many tissues, but short-term fasting reduced CCL2 levels mainly in the liver, spleen and BM among the tissues tested (Figures 4E, S4C and S4D). Strikingly, we found that fasting-induced CCL2 reduction in blood and tissues was lost in *Ppara*^−/−^ mice (Figures 4A, 4B and 4F), and phenformin treatment failed to reduce systemic CCL2 levels in PPARα-deficient animals (Figure 4C).

Hepatocyte-specific deletion of CCL2 neither affected blood CCL2 levels nor the number of blood monocytes (Figure S4E), as was previously shown (Shi et al., 2011). Similarly, splenectomy did not significantly reduce serum CCL2 levels and did not decrease blood monocyte numbers (Figure S4F), suggesting that hepatocytes and splenocytes were not major CCL2 producers for BM monocyte egress.

Previous published data suggested that CCL2 production by BM stromal cells controlled steady-state monocyte egress (Shi et al., 2011). Accordingly, we found that BM CCL2 production correlated with peripheral monocyte numbers (Figure 4G). Immunofluorescence staining of BM tissue sections revealed that CCL2 was produced by cells in close proximity to BM sinusoids (Figure S4G), as was previously suggested (Shi et al., 2011). Also, BM CCL2 production correlated with CCL2 levels found in blood (Figure S4H), suggesting that systemic CCL2 levels might depend on BM CCL2 production. In summary, these data indicated that fasting-induced reduction of BM CCL2 production inhibited monocyte egress to the periphery.

We next examined how fasting-driven hepatic PPARα activation could regulate CCL2 production at distant sites. To this aim, we used mass spectrometry to measure the serum proteome during the fed and the fasting state. We found that fasting led to significant changes in serum protein levels that overlapped with differential gene expression in the fasting liver (Figure 4H and S4I), highlighting the key role of the liver in controlling the composition of the blood plasma proteome in response to dietary intake. We then mined fasting-regulated liver gene transcripts for factors that are released into the circulation and known to modulate CCL2 production. We identified 33 known CCL2 regulators in the liver secretome that were modulated in the fasting state (Figure 4I). Hence, these data suggest that the liver secretome is modified in response to dietary intake and contains multiple circulating factors that are known to regulate CCL2 production, thereby controlling monocyte egress to the periphery.

### Fasting Modifies Monocyte Metabolic Activity

In order to investigate the effect of fasting on monocyte gene expression we profiled monocytes from BM of fed and fasted mice (Figures 5A and 5B). Fasting had a profound effect on the monocyte gene expression profile with more than 2700 genes being differentially expressed. We compared the transcriptional changes found in monocytes from fasting mice to the profile of monocytes from *Ccr2*-deficient mice that cannot sense CCL2 and therefore are unable to mobilize from BM (Serbina and Pamer, 2006). Of note, the transcriptional changes in monocytes from fasting mice overlapped with those of *Ccr2*-deficient monocytes emphasizing the key role of CCL2 in the regulation of monocyte egress upon fasting (Figures 5C and 5D).

**Figure 5.**
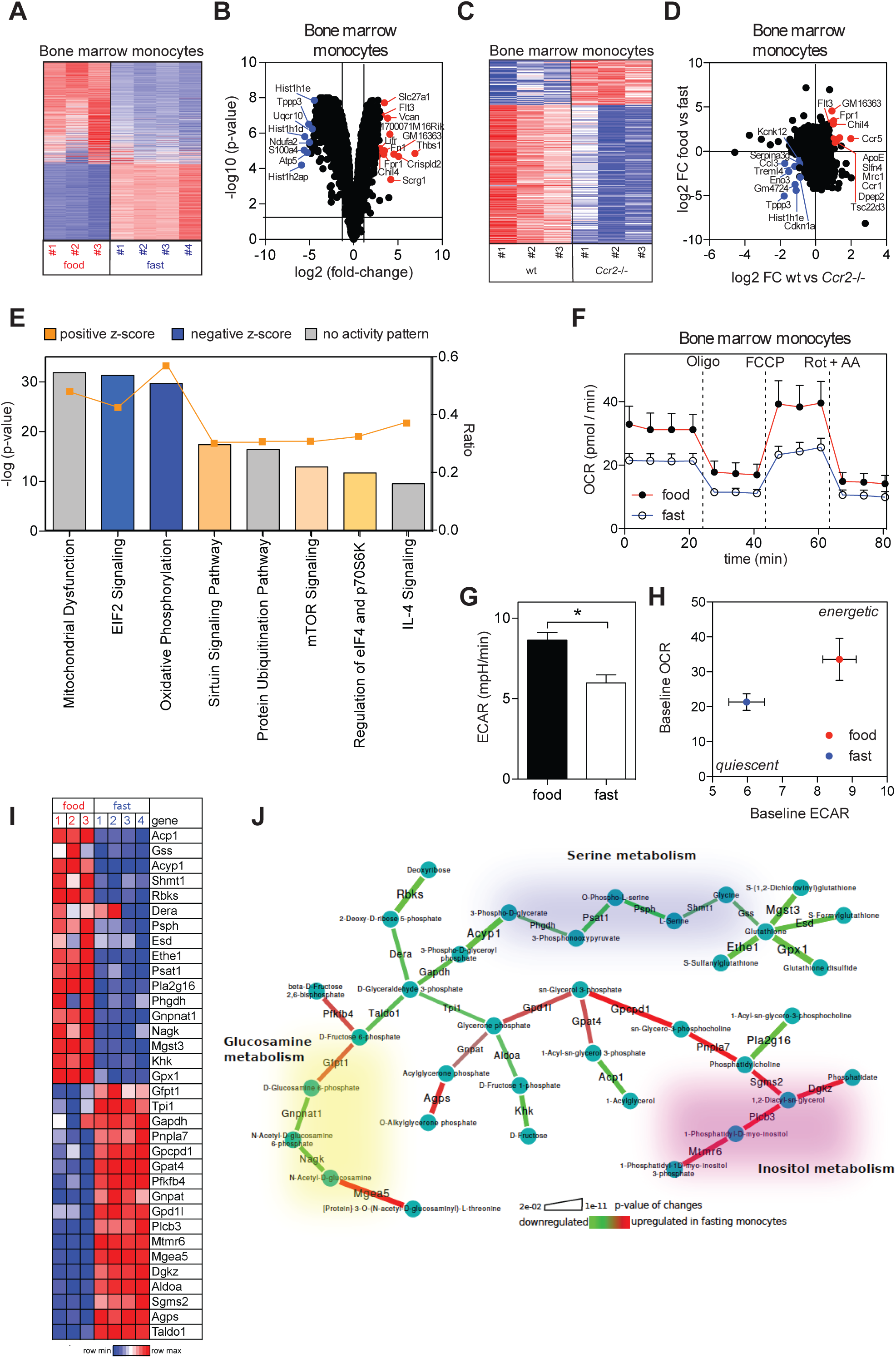
Fasting Modifies Monocyte Metabolic Activity. (A) Heatmap displays z-scores for differentially expressed genes in monocytes purified from the bone marrow of fed and fasted mice. (B) Volcano plot shows top up- and down-regulated transcripts in monocytes from fasted mice compared to monocytes from fed mice. (C) Heatmap displays z-scores for differentially expressed genes in monocytes from bone marrow of wt and *Ccr2*^−/−^ mice. (D) Differentially expressed genes in monocytes between wt and *Ccr2*^−/−^ mice were mapped on differentially expressed genes from monocytes between fed and fasted mice. (E) Ingenuity pathway analysis (IPA) of differentially expressed genes in monocytes between fed and fasted mice. (F) Oxygen consumption rate (OCR) of bone marrow monocytes from fed and fasted mice. Oligo = oligomycin, inhibits ATP-synthase; FCCP = carbonyl cyanide-4 (trifluoromethoxy)phenylhydrazone, mitochondrial uncoupler; Rot + AA = rotenone + antimycin A, CI and CIII inhibitors, respectively. Vertical bars = SEM. (G) Basal extracellular acidification rate (ECAR) of bone marrow monocytes from fed and fasted mice. Vertical bars = SEM. (H) Basal OCR vs. basal ECAR (mean +/-SEM for both parameters). (I and J) Integrated metabolic network analysis of the transcriptional differences between fed and fasting conditions. (I) Heatmap shows scores for subnetwork metabolic enzymes. (J) Graphical representation of the regulated metabolic subnetwork.

Consistent with cellular energy conservation, fasting mostly affected pathways involved in eIF2 signaling and protein ubiquitination as well as mitochondrial function and oxidative phosphorylation (Figure 5E). In fact, monocytes isolated from fasting mice displayed reduced basal (untreated) and maximal (FCCP-induced) oxygen consumption rates (OCR) (Figure 5F) and a reduced basal extracellular acidification (ECAR) rate (Figure 5G), resulting in a quiescent metabolic phenotype compared to monocytes from fed mice (Figure 5H). In depth metabolic network analysis revealed up-regulation of inositol-triphosphate metabolism, which could be indicative of a specific signaling axis during fasting (Figures 5I and 5J). Also, we observed coordinated suppression of serine and glutathione metabolism in monocytes from fasting mice, contrary to what was described for immune cell proliferation upon activation (Ma et al., 2017). Hence, monocytes from fasting mice were reduced in their metabolic activity reflecting a quiescent functional state.

### Fasting Improves Chronic Inflammatory Diseases Without Compromising Monocyte Emergency Mobilization During Acute Inflammation

Caloric restriction improves chronic inflammatory and autoimmune disorders, such as multiple sclerosis and rheumatoid arthritis (Choi et al., 2017; Piccio et al., 2008). Intriguingly, gene modules associated with “inflammation of joint” and “rheumatoid arthritis” were reduced in monocytes isolated from fasting mice (Figure S5A and S5B). Thus, we asked whether fasting-induced changes in monocyte molecular program could improve chronic inflammatory disease outcome. We focused our analysis on mice induced to develop experimental autoimmune encephalomyelitis (EAE), the main preclinical model for multiple sclerosis. EAE progression is strongly dependent on the recruitment of monocytes to the central nervous system, and *Ccr2-*deficient mice, in which monocytes are unable to mobilize from BM, are resistant to disease induction (Ikizler et al., 2018; King et al., 2009; Mildner et al., 2009). Accordingly, disease progression to the paralytic stage correlates with the magnitude of the myeloid infiltrate (Ajami et al., 2011). Strikingly, we observed a strong reduction in myeloid cell accumulation in the spinal cord from fasted mice with EAE (Figure 6A), and intermittent fasting significantly ameliorated EAE clinical course (Figure 6B), as previously shown (Cignarella et al., 2018). Accordingly, gene modules related to infiltration and pro-inflammatory activity were strongly down-regulated in monocytes from fasting mice compared to non-fasting mice (Figure 6C). These data suggest that fasting reduces myeloid cell accumulation in lesional sites, which contributes to improved disease outcome in EAE.

**Figure 6.**
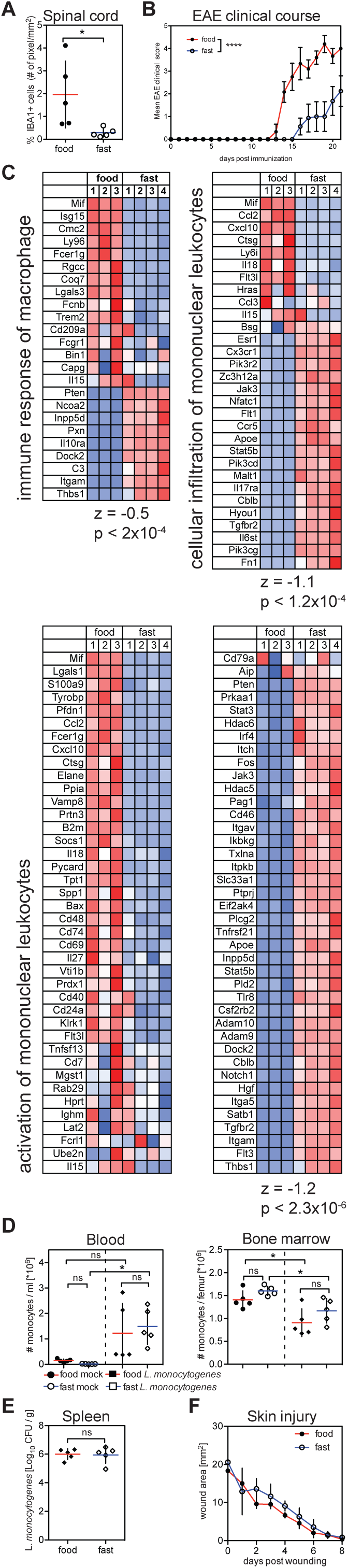
Fasting Improves Chronic Inflammatory Disease Outcome Without Compromising Monocyte Emergency Mobilization During Acute Inflammation. (A and B) Proportion of IBA1+ myeloid cells in spinal cords and (A) and EAE clinical course (B) of mice that were induced with EAE and fed *ad libitum* (food) or underwent intermittent fasting in 24hr cycles (fast). (C) Differentially expressed genes in monocytes between fed and fasted mice were analyzed using ingenuity pathway analysis (IPA). Heatmaps show z-scores of genes related to the indicated gene sets. (D) Absolute numbers of Ly-6C^high^ monocytes in the blood and bone marrow from fed and fasted mice that were infected with *L. monocytogenes*. (E) Colony-forming units (CFU) in the spleen from fed and fasted mice that were infected with *L. monocytogenes*. (F) Wound healing kinetics in mice that were fed *ad libitum* or underwent intermittent fasting. (A, D, E) Every dot represents one individual animal. Horizontal bar = mean. Vertical bar = SD. Two-way analysis of variance (ANOVA) (B, F) or One-way ANOVA with Bonferroni’s post test (D) or Student’s t test (A, E) were performed. Statistical significance is indicated by *P < 0.05, **P < 0.01, ****P < 0.0001. ns = not significant. See also Figure S5.

These results prompted us to ask whether fasting-induced inhibition of monocyte BM egress as well as changes in monocyte functional state would also compromise acute inflammatory reactions in response to tissue injury or pathogen invasion. Both tissue restoration after injury and therapeutic immunity against *Listeria monocytogenes* critically depend on monocytes and are strongly compromised in *Ccr2*-deficient mice (Boniakowski et al., 2018; Serbina and Pamer, 2006). Most interestingly, we found that emergency mobilization upon *Listeria monocytogenes* infection remained intact during fasting (Figure 6D). In addition, neither immunity against *Listeria monocytogenes* nor wound healing were impaired in fasted mice when compared to fed mice (Figure 6E and 6F), suggesting that monocytes accumulating at the site of acute infection or tissue injury remained functional.

In summary, we found that fasting improved chronic inflammatory disease outcome without compromising monocyte emergency mobilization and functionality during acute inflammatory responses.

## DISCUSSION

Mass cytometry profiling of blood cells isolated from healthy humans and mice during fasting revealed a drastic effect of caloric restriction on the blood and tissue monocyte pool. Here we show that dietary intake controls the metabolic and inflammatory activity of BM monocytes and their egress to the blood circulation.

Egress of BM monocytes to the blood circulation is a critical step in the inflammatory cascade induced upon tissue injury (Shi and Pamer, 2011). Monocytes are important producers of pro-inflammatory cytokines and play a critical role in the induction and maintenance of inflammation. Therefore, it is conceivable that modulation of peripheral monocyte load in blood and tissues by repetitive short-term fasting or caloric restriction decreases susceptibility to pathological inflammatory disease. Indeed, circulating monocytes are increased in overweight and obese humans and caloric restriction has been shown to reduce peripheral pro-inflammatory cells leading to an overall improved inflammatory profile (Kani et al., 2017; Poitou et al., 2011). We found that fasting not only reduced the size of the circulating monocyte pool but also modified metabolic activity and gene expression patterns predicting improvement of chronic inflammatory and autoimmune disorders, such as rheumatoid arthritis and multiple sclerosis, diseases that were shown to be responsive to fasting in clinical trials (Choi et al., 2017; Darlington et al., 1986; Kjeldsen-Kragh et al., 1991; Skoldstam et al., 1979). Accordingly, caloric restriction and intermittent fasting strongly reduced the accumulation of pathogenic monocytes in the central nervous system and ameliorated disease outcome in a preclinical model of multiple sclerosis (Cignarella et al., 2018; Piccio et al., 2008).

While decreased inflammatory activity is desirable in chronic autoimmune diseases, it might be devastating in response to tissue injury or pathogen invasion. Importantly, we found that fasting did not compromise tissue regeneration nor immunity against *Listeria monocytogenes*, a condition most critically dependent on monocyte emergency mobilization for the induction of therapeutic immunity. Fasting as part of physiological sickness behavior can even be beneficial in antimicrobial immunity (Wang et al., 2016).

It has been speculated for a long time that hepatocytes might modulate monocyte BM hematopoiesis and egress (Decker et al., 2018; Serbina et al., 2012). Here we show that food energy-sensing by a liver AMPK-PPARα pathway controls BM monocyte egress to the blood circulation. Interestingly, anti-inflammatory dietary patterns such as the Mediterranean diet are highly enriched in natural PPARα agonists. In fact, PPARα-dependent fasting metabolites are associated with anti-inflammatory effects (Wang et al., 2016). β-Hydroxybutyrate, for example, exerts anti-inflammatory functions through binding to GPR109A or by inhibiting the NLRP3 inflammasome (Yamanashi et al., 2017; Youm et al., 2015). We have found that hepatic AMPK and PPARα not only coordinated metabolic adaptation to fasting, but also controlled the pool of circulating inflammatory cells through the modulation of steady-state CCL2 production.

Caloric restriction improves inflammatory and autoimmune disease outcome. Importantly, our finding that pharmacological AMPK activation regardless of caloric intake was sufficient to regulate the blood monocyte pool suggests that targeting liver energy sensors could be an innovative strategy for the prevention and treatment of chronic inflammatory and autoimmune diseases without affecting antimicrobial immunity.

Altogether, these data establish diet composition and liver energy sensors as critical regulators of the blood and tissue inflammatory tone and identify novel clinical strategies for the treatment of patients with chronic inflammatory diseases and autoimmune disorders.

## ACKNOWLEDGEMENTS

We thank J. Agudo, S. Offermanns, C. Buettner, S. Fried, T. E. McGraw, and the Merad laboratory for helpful discussions. We thank A. Lansky for help with the IRB application, the Flow Cytometry facility for technical support, the Human immune monitoring core for assistance with the multiplex assay, M. Davila for processing CyTOF data, M. Serasinghe for help with metabolic measurements in monocytes, the Stable Isotope & Metabolomics core at Albert Einstein College of Medicine for assistance with blood metabolite measurements, S. Hatem for assistance with phlebotomy and all participants of the human fasting experiment. We are grateful to Y. Belkaid for giving us the opportunity to use her laboratory at NIH and N. Bouladoux for assistance. Supported by the US National Institutes of Health (to M.M., R01 NS102633-01 to L.P., K08CA190770 to E.J.G), German Research Council (DFG) (SFB-TRR57 P07 to M.-L.B, JO 1216/1-1 to S.J.)

## AUTOR CONTRIBUTIONS

S.J. conceived the study, designed, performed and analyzed experiments and wrote the manuscript. N.T., M.C.A., C.C., Cl. C., D.Z., T.H.W., S.N., B.B.M., A.R. and F.M. performed experiments and analyzed data. S.A.R., A.G. and M.N.A. analyzed the RNA-seq data. J.E.C., D.L. and P.S.F. provided intellectual expertise. C.N.B. and F.J.G. provided mice and intellectual expertise. L.P. and M.-L.B. designed and funded experiments. E.J.G. performed, analyzed and funded experiments and wrote the manuscript. M.M. supervised and funded the study and wrote the manuscript. All authors discussed the data and edited the manuscript.

## STAR METHODS

### EXPERIMENTAL MODEL AND SUBJECT DETAILS

#### Animals

C57BL/6 were purchased from Charles River at the age of 6 weeks and housed in our facility for at least two weeks before being used in experiments. B6;129S4-*Ppara*^*tm1Gonz*^/J (*Ppara*^−/−^ mice; Stock No.: 008154) were purchased from Jackson and bred in our facility. B6.129P2-*Lyz2*^tm1(cre)Ifo^/J (LysMcre; 004781) and B6.129S4(FVB)-*Insr*^tm1Khn^/J (*Insr*^fl/fl^ mice; 006955) mice were crossed to delete the *Insr* gene from monocytes (*Insr*^**Δ**Mono^ mice). B6.Cg-*Speer6*-^ps1Tg(Alb-cre)21Mgn^/J (*Alb*^cre/cre^; 003574) mice were crossed to *Prkaa1*^tm1.1Sjm^/J (AMPK^fl/fl^; 014141), B6.129S6(SJL)-*Fgf21*^tm1.20jm^/J (*Fgf21*^fl/fl^; 022361) and B6.Cg-*Ccl2*^tm1.1Pame^/J (*Ccl2*^fl/fl^; 016849) mice in order to generate hepatocyte-specific deletions of AMPK, *Fgf21* and *Ccl2* (AMPK^**Δ**Hep^, *Fgf21*^**Δ**Hep^, *Ccl2*^**Δ**Hep^). For experiments involving cre-positive conditional knockout mice, cre-negative littermates were used as controls. Mice were housed at specified pathogen free (SPF) health status in individually ventilated cages at 21-22°C and 39-50% humidity in groups of 5 animals. Female mice at the age of 8-12 weeks were used for the experiments. Male conditional *Ppara*^ΔHep^ mice were described earlier (Brocker et al., 2017) and housed at the National Cancer Institute. Intermittent fasting experiments in mice induced with EAE were performed at Washington University in St. Louis. All animal procedures performed in this study were approved by the Institutional Animal Care and Use Committee (IACUC) of the respective institutions.

#### Human fasting trial

Healthy individuals (3 female and 9 male participants, mean age = 30+/-5 years) with a BMI 18.5-25 kg/m^2^ were admitted to the study. Leukocyte populations in human blood during the eating and the fasting state were assessed using a 2-day protocol. On the first day participants ate before 12 pm and did not consume food until after 3 pm (Figure 1A). At 3 pm blood was drawn by venipuncture (eating state) into an EDTA-coated tube. After the first blood draw participants were allowed to consume food until 8pm. Participants fasted from 8 pm until 3 pm of the following day, when the second blood sample was obtained (fasting state). Participants were allowed to drink water at all times. On both days, the total white blood count was measured using a hemocytometer (Becton Dickinson). Cells were processed for CyTOF immediately after the blood draw. The study was approved by the institutional review board and informed consent was obtained from all subjects.

#### Infection with *Listeria monocytogenes*

*Listeria monocytogenes* serovar 1/2a EGDe was a kind gift from Werner Göbel, Max von Pettenkofer-Institute, Ludwig-Maximilians-Universität München. Bacteria were grown in liquid BBL Brain Heart Infusion (BHI) medium (211059, BD) at 37°C until the culture reached OD_600_=1. Cells were washed twice in icecold PBS, resuspended and frozen in aliquots in PBS 20% glycerol. Infectious units were determined by titration on BHI agar plates (255003, BD) Mice were infected intravenously with 3000 CFU and organs were harvested 24 hrs later. Spleen homogenates were plated on BHI agar plates.

### METHOD DETAILS

#### Diets

Mice were fed irradiated rodent Diet 20 (5053, PicoLab) *ad libitum* unless indicated otherwise and had access to reverse osmosis water from autoclaved bottles or an automatic watering system. For measurements of food intake purified diet AIN-93G (TD.94045, Envigo) was used. An AIN-93G matching fasting diet devoid of macronutrients was prepared as described before (Brandhorst et al., 2013). In brief, 0.43 ml essential fatty acids (Udo’s Oil 3*6*9, Whole Foods), 10 g fiber (Cellulose #3425, Bio-Serv), 7 g AIN-93G-MX (TD.94046, Envigo), 2 g AIN-93-VX (TD.94047, Envigo) were mixed with 180.60 g hot hydrogel (70-01-5022, Clear H2O). AIN93-G intake of individual mice was measured every day for a week and the amount of fasting diet fed every 24 hrs was adjusted to the previously measured 24 hr food intake. For ketogenic diet experiments ketogenic diet F3666 (BioServ) and control diet AIN-93M (TD.94048, Envigo) were used.

#### Experimental animal fasting

For 4 hr fasting protocols, food was removed at 9 am (ZT2), and for overnight fasting food was removed at 5 pm (ZT10) and the cage bedding and nesting material were changed to prevent coprophagy. Control mice were fed *ad libitum*. Fasting and eating mice had access to water *ad libitum*.

#### Treatments

For β3-AR stimulation, mice were gavaged with 1 mg/kg CL 316,243 (C5976, Sigma) dissolved in water 3 hrs before analysis. For gavage of macronutrients 0.2 ml of 30% glucose (w/v) (G8270, Sigma), 30% BSA (w/v) (BAH62, Equitech-Bio) and 15% extra virgin olive oil (v/v) (Bertolli) was used as described before (Wang et al., 2016). Mice were gavaged once every hour for 4 hrs. For hexokinase inhibition mice were gavaged with 750 mg/kg 2-deoxyglucose (D6134, Sigma) or 170 mg/kg D-mannoheptulose (97318, Sigma) in water once every hour for 4 hrs. For pharmacological AMPK activation mice were gavaged with a single dose of 50 mg/kg phenformin HCl (S2542, Selleckchem) dissolved in water unless indicated otherwise, or 50 mg/kg A-769662 (S2697, Selleckhem) suspended in 1% carboxymethylcellulose (C4888, Sigma), and blood and bone marrow were analyzed 4 hrs later. Insulin (HumulinR, Lilly) was injected intraperitoneally into mice that fasted for 3 hrs at a dose of 1 unit/kg. For CCL2 reconstitution mice were fasted overnight and received 80 ug/kg CCL2 protein (479-JE/CF, R&D) in PBS intravenously 3 hrs before analysis. 4hr treatments were started at 9am (ZT2), 3hr treatments at 10am (ZT3) in the home cage and mice were analyzed at 1pm (ZT6).

#### Induction of experimental autoimmune encephalomyelitis (EAE)

Mice were fasted in 24 hr cycles for 4 weeks before induction of EAE (Cignarella et al., 2018). For disease induction, a commercially available kit was used according to the manufaturer’s instructions (EK-2110, Hooke Laboratories).

#### Punch biopsy

The dorsal skin of mice was shaved and 6 mm biopsy punches (Miltex) were used to make full-thickness wounds. Wound closure was assessed macroscopically with an engineer’s caliper daily. Mice were fasted in 24 hr cycles during the experiment.

#### Bone marrow chimeras

Bone marrow chimeras were generated by transplantation of 2×10^6^ bone marrow donor cells into lethally irradiated (2×6.5Gy) recipient mice. Mice were kept on sulfamethoxazole / trimethoprim (STI Pharma) for 3 weeks and analyzed 7 weeks after the transfer.

#### Flow cytometry

Unless indicated otherwise organs were harvested at 1 pm (ZT6). Single cell suspensions were obtained from spleens by passing the organ through a 100 um cell strainer and from liver and lung after digestion with 0.5mg / ml collagenase IV (C5138, Sigma) at 37°C for 45 and 30 min, respectively. For liver tissue, nonparenchymal cells were enriched by centrifugation in 35% Percoll (17-0891-01, GE Healthcare) for 30 min at 1,300 rcf. Single cell suspension from adipose tissue was prepared as described before (Orr et al., 2013). In brief, gonadal fat pads were cut into small pieces and digested in collagenase IV 10 mM CaCl_2_ at 37°C on a shaker at 100 rpm for 20 min. Digested tissue was triturated and passed through a 100 um filter. Liver, lung, adipose tissue, spleen and bone marrow single cell suspensions were incubated with ACK lysing buffer (420301, BioLegend) for 3 min at room temperature. Blood was drawn retro-orbitally or from liver sinus into EDTA-coated MiniCollect tubes (450475, Greiner). 50 ul blood were transferred into 2 ml FACS buffer (PBS w/o Ca^2+^ and Mg^2+^ supplemented with 2% heat inactivated FBS and 5 mM EDTA) and treated twice with ACK lysing buffer for 5 min at room temperature. For flow cytometry, cells were stained in FACS buffer with mAbs specific to CD45 (clone 30-F11, BioLegend), CD45.1 (A20, eBioscience), CD45.2 (104, eBioscience), CD11b (M1/70, eBioscience), Ly-6G (1A8, BioLegend), Ly-6C (AL-21, BD Biosciences), CD115 (AFS98, eBioscience), CX3CR1 (SA011F11, BioLegend), CXCR4 (2B11, eBioscience) for 20 min on ice. Proapoptotic cells were stained for Caspase-3/7 activation using the Vybrant FAM Caspase-3 and -7 assay kit (V35118, Invitrogen) following the manufacturer’s instructions. Dead cells were excluded using DAPI staining (D1306, Life technologies) and AccuCheck counting beads (PCB100, Molecular Probes) were used for quantification of absolute cell numbers. Multiparameter analysis was performed on an LSR Fortessa II (Becton Dickinson) using the FACSDiva software and data were analyzed using the Flo Jo software (Tree Star Inc.).

#### Cytometry by time-of-flight spectrometry (CyTOF)

All mass cytometry reagents were purchased from Fluidigm Inc. unless otherwise noted. Individual human whole blood samples were stained with a cocktail of the following metal labeled antibodies (all antibodies sourced from Biolegend and conjugated in house using Fluidigm X8 MaxPar conjugation kits unless otherwise noted): CD11c 115 In (Bu15), CD33 141 Pr (WM53), CD19 142 Nd (HIB19), CD45RA 143 Nd (HI100), CD141 144 Nd (M80), CD4 145 Nd (RPA-T4), CD8 146 Nd (RPA-T8), CD370 147 Sm (8F9), CD16 148 Nd (3G8), CD1c 150 Nd (L161), CD123 151 Eu (6H6), CD66b 152 Sm (G10F5), CCR2 153 Eu (K036C2; Fluidigm), CD86 154 Sm (IT2.2), CD27 155 Gd (O323), PDL1 156 Gd (29E.2A3), CD163 158 Gd (GHI/61), CD24 159 Tb (ML5), CD14 160 Gd (M5E2), CD56 161 Dy (B159; BD Biosciences), CD64 162 Dy (10.1), CD172a/b 163 Dy (SE5A5), CD40 164 Dy (5C3), CCR6 165 Ho (G034E3), CD25 166 Er (M-A251), CCR7 167 Er (G043H7), CD3 168 Er (UCHT1), CX3CR1 169 Tm (2A9-1), CD38 170 Er (HB-7), CD161 171 Yb (HP-3G10), CXCR3 173 Yb (G025H7), HLADR 174 Yb (L243), Axl 175 Lu (108724; R&D), CCR4 176 Yb (205410; R&D), CD11b 209 Bi (M1/70). The titrated panel of antibodies was added directly to 400 ul of whole blood and incubated for 20 min at RT. The blood was then fixed and lysed using BD FACS Lysing solution, and the cells were post-fixed for 30 min with freshly diluted 1.6% formaldehyde in PBS containing a 1:4000 dilution of Ir nucleic acid intercalator to label all nucleated cells. Staining cells were stored in PBS containing 0.1% BSA until immediately prior to acquisition.

Individual mouse blood samples were barcoded with anti CD45 antibodies (clone A20) and pooled for batched analysis. Samples were stained with a cocktail of the following metal-conjugated antibodies from Biolegend unless noted otherwise: Ly-6G 141 Pr (clone 1A8), CD11c 142 Nd (N418), TCRb 143 Nd (H57-597), CD8 168 Er (53-6.7), CD11b 148 Nd (M1/70), CD19 149 Sm (6D5), CD24 144 Nd (M1/69), CD25 151 Eu (3C7), Siglec-F 152 Sm (E50-2440; BD Bioscience), CD64 156 Gd (X54-5/7.1), NK1.1 170 Er (PK136), CD62L 160 Gd (MEL-14), Ly-6C 162 Dy (HK1.4), CD103 161 Dy (2E7), CD117 166 Er (2B8), CD44 171 Yb (IM7), CD4 172 Yb (RM4-5), MHCII 174 Yb (M5/114.15.2), Thy1.2 113 In (30-H12), F4/80 146 Nd (BM8), IgD 150 Nd (11-26c.2a), CD169 154 Sm (3D6.112), CX3CR1 176 Yb (SA011F11), Cisplatin 195 Pt.

The cells were stained for 30 min on ice, and then washed. After antibody staining, the cells were incubated with cisplatin for 5 min at RT as a viability dye for dead cell exclusion. Cells were fixed in PBS containing 1.6% formaldehyde and a 1:4000 dilution of Ir nucleic acid intercalator to label all nucleated cells.

Immediately prior to acquisition, the cells were washed in PBS, then in diH20 and resuspended in diH_2_0 containing a 1/10 dilution of Equation 4 Element Calibration beads. After routine instrument tuning and optimization, the samples were acquired on a CyTOF2 Mass Cytometer equipped with a Super Sampler fluidics system (Victorian Airships) at an acquisition rate of < 500 events /s. The resulting FCS files were concatenated and normalized using a bead-based normalization algorithm in the CyTOF acquisition software and uploaded to Cytobank for analysis. FCS files were manually pre-gated on Ir193 DNA+ CD45+ events, excluding dead cells, doublets and DNA-negative debris, and the gated populations were then analyzed using viSNE (Amir el et al., 2013) and Phenograph.

#### Hepatocyte isolation

For primary hepatocyte isolation the inferior vena cava was cannulated and livers were perfused with calcium-free salt solution containing 0.5 mM EGTA followed by perfusion with 90 ml of a 0.02% collagenase D solution (11088882, Roche) at RT. After digestion livers were gently minced on a Petri dish and passed through a 70 um cell strainer. Hepatocytes were washed three times in PBS and centrifuged at 50xg for 3 min. Cells were resuspended in Trizol at 200,000 cells / ml for further analysis.

#### ELISA and multiplex

Blood serum was isolated using Z-Gel 1.1 ml serum preparation tubes (41.1378.005, Sarstedt) according to the manufacturer’s instructions. Tissue for protein measurements was snap frozen in liquid nitrogen. 50-100 mg tissue were added to 10x the weight T-PER Tissue Protein Extraction Reagent (78510, ThermoFisher) containing Halt Proteinase and Phosphatase inhibitor cocktail (78440, ThermoFisher). Tissue was homogenized using a 5 mm steel bead (Qiagen) and a TissueLyser II (Qiagen) at 30 hz for 2 min (spleen, liver, adipose tissue, pancreas, thymus, kidney), 5 min (muscle, heart, lung, cecum, large intestine, small intestine) or 10 min (skin, stomach). Debris was pelleted and supernatant used for assays. Bone marrow was prepared by flushing 2 femurs with 500 ul PBS. After pelleting the cells, supernatant was used for BMEF measurements, while cells were taken up in T-PER lysis buffer, underwent one freeze-thaw cycle and debris was pelleted. For measurement of serum and tissue chemokine levels commercially available ELISA kits for CCL2/JE (MJE00, R&D) and CXCL12/SF-1 (MCX120, R&D) were used. For measurement of blood stress hormone corticosterone an ELISA kit was used (ADI-900-097, Enzo). Multiplex was performed using a kit for mouse metabolic hormones (MMHMAG-44K, Millipore) and human metabolic hormones (HMEMAG-34K, Millipore).

#### Serum metabolomics

Commercially available kits were used to measure total serum triglycerides (T7532, Pointe Scientific) and β-hydroxybutyrate (700190, Cayman chemicals) according to the manufacturer’s instructions. Measurements for metabolites in mouse plasma were performed using the AbsoluteIDQ p180 kit (Biocrates) by the Stable Isotope & Metabolomics Core at Albert Einstein College of Medicine, New York.

#### qPCR

Conventional reverse transcription was performed using the RNA to cDNA EcoDry Premix (ST0335, Clontech) in accordance with the manufacturer’s instructions. qPCR was performed on a CFX384 Touch Real-Time PCR detection system (Bio Rad) using EXPRESS SYBR GreenER master mix (11784200, Invitrogen) and primers for *Ppara* (5’-*Ppara* AGAGCCCCATCTGTCCTCTC, 3’-*Ppara* ACTGGTAGTCTGCAAAACCAAA) and *ActB* (5’-*ActB* TTCCTTCTTGGGTATGGAATCCTG, 3’-*ActB* GAGGTCTTTACGGATGTCAACG) as follows: one cycle at 95°C (10 min), 40 cycles of 95°C (15 s) and 58°C (1 min). Expression of *ActB* was used as a standard. The average threshold cycle number (CRtR) for each tested mRNA was used to quantify the relative expression: 2^[Ct(*Actb*)-Ct(*Ppara*)].

#### Metabolic measurements in monocytes

For metabolic measurements 250.000 monocytes / well were plated in a XF96 plate. We used the Seahorse XF Cell Mito Stress Test Kit (103015-100, Agilent) according to the manufacturer’s instructions.

#### Transcriptome analysis

Transcript abundances were quantified using the Ensembl GRCm38 cDNA reference using Kallisto version 0.44.0. Transcript abundances were summarized to gene level using tximport. Expression matrices were filtered for only transcripts with greater than 5 TPM in all replicates of at least one condition. Differential expression statistics between different conditions were calculated using limma/voom method in R with TMM normalization. P values were adjusted for multiple testing by Benjamini-Hochberg correction.

#### Bioinformatics

For bioinformatical analysis of transcriptional profiling data we used the ingenuity pathway analysis portal (Qiagen). For identification of potential metabolic modules differentially regulated in monocytes during the fed and fasting state a method called GATOM (from Genes, Metabolites and Atoms) was used (Sergushichev et al., 2016; Ulland et al., 2017). GATOM uses carbon atomtransition graph. It is a graph where each vertex is an atom of a metabolite and an edge connects two atoms A1 and A2 of metabolites M1 and M2 if there is a reaction R with M1 and M2 on different sides of equation and atom A1 of M1 transforms to atom A2 of M2. Each element (vertex or edge) of the graph is assigned with a weight that is positive if the data support its importance or negative if not. The weight could be derived from statistical test p-values, such as differential expression, carried out for transcriptional or metabolic data.

### QUANTIFICATION AND STATISTICAL ANALYSIS

#### Statistics

Unless indicated otherwise, data are shown from one out of at least two experiments with n > 3 mice per group. Data are plotted for individual animals with group means (horizontal lines) and standard deviation (SD, vertical lines). Statistical analysis was performed using Prism 5 (GraphPad Software). Depending on the dataset two-tailed Student’s t test or TWO- or ONE-way ANOVA with Tukey’s, Dunnett’s or Bonferroni’s multiple comparison tests were applied to determine significance, and the test used for statistical analysis is indicated in the figure legend. Statistical significance is indicated by *P < 0.05, **P < 0.01, ***P < 0.001, ****P < 0.0001. ns = not significant.

## REFERENCES

Ajami, B., Bennett, J.L., Krieger, C., McNagny, K.M., and Rossi, F.M. (2011). Infiltrating monocytes trigger EAE progression, but do not contribute to the resident microglia pool. Nat Neurosci 14, 1142– 1149.

Aksungar, F.B., Topkaya, A.E., and Akyildiz, M. (2007). Interleukin-6, C-reactive protein and biochemical parameters during prolonged intermittent fasting. Ann Nutr Metab 51, 88–95.

Boniakowski, A.E., Kimball, A.S., Joshi, A., Schaller, M., Davis, F.M., denDekker, A., Obi, A.T., Moore, B.B., Kunkel, S.L., and Gallagher, K.A. (2018). Murine macrophage chemokine receptor CCR2 plays a crucial role in macrophage recruitment and regulated inflammation in wound healing. European journal of immunology 48, 1445–1455.

Brocker, C.N., Yue, J., Kim, D., Qu, A., Bonzo, J.A., and Gonzalez, F.J. (2017). Hepatocyte-specific PPARA expression exclusively promotes agonist-induced cell proliferation without influence from nonparenchymal cells. American journal of physiology Gastrointestinal and liver physiology 312, G283–G299.

Cheng, C.W., Villani, V., Buono, R., Wei, M., Kumar, S., Yilmaz, O.H., Cohen, P., Sneddon, J.B., Perin, L., and Longo, V.D. (2017). Fasting-Mimicking Diet Promotes Ngn3-Driven beta-Cell Regeneration to Reverse Diabetes. Cell 168, 775–788 e712.

Choi, I.Y., Lee, C., and Longo, V.D. (2017). Nutrition and fasting mimicking diets in the prevention and treatment of autoimmune diseases and immunosenescence. Mol Cell Endocrinol 455, 4–12.

Choi, I.Y., Piccio, L., Childress, P., Bollman, B., Ghosh, A., Brandhorst, S., Suarez, J., Michalsen, A., Cross, A.H., Morgan, T.E., etal. (2016). A Diet Mimicking Fasting Promotes Regeneration and Reduces Autoimmunity and Multiple Sclerosis Symptoms. Cell Rep 15, 2136–2146.

Chong, S.Z., Evrard, M., Devi, S., Chen, J., Lim, J.Y., See, P., Zhang, Y., Adrover, J.M., Lee, B., Tan, L., et al. (2016). CXCR4 identifies transitional bone marrow premonocytes that replenish the mature monocyte pool for peripheral responses. The Journal of experimental medicine 213, 2293–2314.

Cignarella, F., Cantoni, C., Ghezzi, L., Salter, A., Dorsett, Y., Chen, L., Phillips, D., Weinstock, G.M., Fontana, L., Cross, A.H., etal. (2018). Intermittent Fasting Confers Protection in CNS Autoimmunity by Altering the Gut Microbiota. Cell metabolism 27, 1222–1235 e1226.

Darlington, L.G., Ramsey, N.W., and Mansfield, J.R. (1986). Placebo-controlled, blind study of dietary manipulation therapy in rheumatoid arthritis. Lancet 1, 236–238.

Decker, M., Leslie, J., Liu, Q., and Ding, L. (2018). Hepatic thrombopoietin is required for bone marrow hematopoietic stem cell maintenance. Science 360, 106–110.

Faris, M.A., Kacimi, S., Al-Kurd, R.A., Fararjeh, M.A., Bustanji, Y.K., Mohammad, M.K., and Salem, M.L. (2012). Intermittent fasting during Ramadan attenuates proinflammatory cytokines and immune cells in healthy subjects. Nutrition research 32, 947–955.

Fontana, L., Partridge, L., and Longo, V.D. (2010). Extending healthy life span-from yeast to humans. Science 328, 321–326.

Haslam, D.W., and James, W.P. (2005). Obesity. Lancet 366, 1197–1209.

Ho, T.P., Zhao, X., Courville, A.B., Linderman, J.D., Smith, S., Sebring, N., Della Valle, D.M., Fitzpatrick, B., Simchowitz, L., and Celi, F.S. (2015). Effects of a 12-month moderate weight loss intervention on insulin sensitivity and inflammation status in nondiabetic overweight and obese subjects. Horm Metab Res 47, 289–296.

Ikizler, T.A., Robinson-Cohen, C., Ellis, C., Headley, S.A.E., Tuttle, K., Wood, R.J., Evans, E.E., Milch, C.M., Moody, K.A., Germain, M., et al. (2018). Metabolic Effects of Diet and Exercise in Patients with Moderate to Severe CKD: A Randomized Clinical Trial. J Am Soc Nephrol 29, 250–259.

Imayama, I., Ulrich, C.M., Alfano, C.M., Wang, C., Xiao, L., Wener, M.H., Campbell, K.L., Duggan, C., Foster-Schubert, K.E., Kong, A., et al. (2012). Effects of a caloric restriction weight loss diet and exercise on inflammatory biomarkers in overweight/obese postmenopausal women: a randomized controlled trial. Cancer research 72, 2314–2326.

Jahromi, S.R., Sahraian, M.A., Ashtari, F., Ayromlou, H., Etemadifar, M., Ghaffarpour, M., Mohammadianinejad, E., Nafissi, S., Nickseresht, A., Shaygannejad, V., et al. (2014). Islamic fasting and multiple sclerosis. BMC Neurol 14, 56.

Jensen, P., Christensen, R., Zachariae, C., Geiker, N.R., Schaadt, B.K., Stender, S., Hansen, P.R., Astrup, A., and Skov, L. (2016). Long-term effects of weight reduction on the severity of psoriasis in a cohort derived from a randomized trial: a prospective observational follow-up study. Am J Clin Nutr 104, 259–265.

Jensen, T.L., Kiersgaard, M.K., Sorensen, D.B., and Mikkelsen, L.F. (2013). Fasting of mice: a review. Lab Anim 47, 225–240.

Johnson, J.B., Summer, W., Cutler, R.G., Martin, B., Hyun, D.H., Dixit, V.D., Pearson, M., Nassar, M., Telljohann, R., Maudsley, S., etal. (2007). Alternate day calorie restriction improves clinical findings and reduces markers of oxidative stress and inflammation in overweight adults with moderate asthma. Free Radic Biol Med 42, 665–674.

Kani, A.H., Alavian, S.M., Esmaillzadeh, A., Adibi, P., Haghighatdoost, F., and Azadbakht, L. (2017). Effects of a Low-Calorie, Low-Carbohydrate Soy Containing Diet on Systemic Inflammation Among Patients with Nonalcoholic Fatty Liver Disease: A Parallel Randomized Clinical Trial. Horm Metab Res 49, 687–692.

King, I.L., Dickendesher, T.L., and Segal, B.M. (2009). Circulating Ly-6C+ myeloid precursors migrate to the CNS and play a pathogenic role during autoimmune demyelinating disease. Blood 113, 3190– 3197.

Kjeldsen-Kragh, J., Haugen, M., Borchgrevink, C.F., Laerum, E., Eek, M., Mowinkel, P., Hovi, K., and Forre, O. (1991). Controlled trial of fasting and one-year vegetarian diet in rheumatoid arthritis. Lancet 338, 899–902.

Lips, M.A., van Klinken, J.B., Pijl, H., Janssen, I., Willems van Dijk, K., Koning, F., and van Harmelen, V. (2016). Weight loss induced by very low calorie diet is associated with a more beneficial systemic inflammatory profile than by Roux-en-Y gastric bypass. Metabolism 65, 1614–1620.

Loria-Kohen, V., Fernandez-Fernandez, C., Bermejo, L.M., Morencos, E., Romero-Moraleda, B., and Gomez-Candela, C. (2013). Effect of different exercise modalities plus a hypocaloric diet on inflammation markers in overweight patients: a randomised trial. Clin Nutr 32, 511–518.

Lumeng, C.N., and Saltiel, A.R. (2011). Inflammatory links between obesity and metabolic disease. The Journal of clinical investigation 121, 2111–2117.

Ma, E.H., Bantug, G., Griss, T., Condotta, S., Johnson, R.M., Samborska, B., Mainolfi, N., Suri, V., Guak, H., Balmer, M.L., et al. (2017). Serine Is an Essential Metabolite for Effector T Cell Expansion. Cell metabolism 25, 482.

Manzel, A., Muller, D.N., Hafler, D.A., Erdman, S.E., Linker, R.A., and Kleinewietfeld, M. (2014). Role of “Western diet” in inflammatory autoimmune diseases. Curr Allergy Asthma Rep 14, 404.

Mendez-Ferrer, S., Lucas, D., Battista, M., and Frenette, P.S. (2008). Haematopoietic stem cell release is regulated by circadian oscillations. Nature 452, 442–447.

Mildner, A., Mack, M., Schmidt, H., Bruck, W., Djukic, M., Zabel, M.D., Hille, A., Priller, J., and Prinz, M. (2009). CCR2+Ly-6Chi monocytes are crucial for the effector phase of autoimmunity in the central nervous system. Brain 132, 2487–2500.

Moro, T., Tinsley, G., Bianco, A., Marcolin, G., Pacelli, Q.F., Battaglia, G., Palma, A., Gentil, P., Neri, M., and Paoli, A. (2016). Effects of eight weeks of time-restricted feeding (16/8) on basal metabolism, maximal strength, body composition, inflammation, and cardiovascular risk factors in resistance-trained males. J Transl Med 14, 290.

Oh, E.G., Bang, S.Y., Kim, S.H., Hyun, S.S., Chu, S.H., Jeon, J.Y., Im, J.A., Lee, J.E., and Lee, M.K. (2013). Therapeutic lifestyle modification program reduces plasma levels of the chemokines CRP and MCP-1 in subjects with metabolic syndrome. Biol Res Nurs 15, 48–55.

Ott, B., Skurk, T., Hastreiter, L., Lagkouvardos, I., Fischer, S., Buttner, J., Kellerer, T., Clavel, T., Rychlik, M., Haller, D., et al. (2017). Effect of caloric restriction on gut permeability, inflammation markers, and fecal microbiota in obese women. Sci Rep 7, 11955.

Picca, A., Pesce, V., and Lezza, A.M.S. (2017). Does eating less make you live longer and better? An update on calorie restriction. Clin Interv Aging 12, 1887–1902.

Piccio, L., Stark, J.L., and Cross, A.H. (2008). Chronic calorie restriction attenuates experimental autoimmune encephalomyelitis. J Leukoc Biol 84, 940–948.

Poitou, C., Dalmas, E., Renovato, M., Benhamo, V., Hajduch, F., Abdennour, M., Kahn, J.F., Veyrie, N., Rizkalla, S., Fridman, W.H., et al. (2011). CD14dimCD16+ and CD14+CD16+ monocytes in obesity and during weight loss: relationships with fat mass and subclinical atherosclerosis. Arterioscler Thromb Vasc Biol 31, 2322–2330.

Ramel, A., Martinez, J.A., Kiely, M., Bandarra, N.M., and Thorsdottir, I. (2010). Effects of weight loss and seafood consumption on inflammation parameters in young, overweight and obese European men and women during 8 weeks of energy restriction. Eur J Clin Nutr 64, 987–993.

Serbina, N.V., and Pamer, E.G. (2006). Monocyte emigration from bone marrow during bacterial infection requires signals mediated by chemokine receptor CCR2. Nature immunology 7, 311–317.

Serbina, N.V., Shi, C., and Pamer, E.G. (2012). Monocyte-mediated immune defense against murine Listeria monocytogenes infection. Adv Immunol 113, 119–134.

Shi, C., Jia, T., Mendez-Ferrer, S., Hohl, T.M., Serbina, N.V., Lipuma, L., Leiner, I., Li, M.O., Frenette, P.S., and Pamer, E.G. (2011). Bone marrow mesenchymal stem and progenitor cells induce monocyte emigration in response to circulating toll-like receptor ligands. Immunity 34, 590–601.

Shi, C., and Pamer, E.G. (2011). Monocyte recruitment during infection and inflammation. Nature reviews Immunology 11, 762–774.

Shibolet, O., Alper, R., Avraham, Y., Berry, E.M., and Ilan, Y. (2002). Immunomodulation of experimental colitis via caloric restriction: role of Nk1.1+ T cells. Clin Immunol 105, 48–56.

Skoldstam, L., Larsson, L., and Lindstrom, F.D. (1979). Effect of fasting and lactovegetarian diet on rheumatoid arthritis. Scand J Rheumatol 8, 249–255.

Starr, M.E., Steele, A.M., Cohen, D.A., and Saito, H. (2016). Short-Term Dietary Restriction Rescues Mice From Lethal Abdominal Sepsis and Endotoxemia and Reduces the Inflammatory/Coagulant Potential of Adipose Tissue. Crit Care Med 44, e509–519.

Tajik, N., Keshavarz, S.A., Masoudkabir, F., Djalali, M., Sadrzadeh-Yeganeh, H.H., Eshraghian, M.R., Chamary, M., Ahmadivand, Z., Yazdani, T., and Javanbakht, M.H. (2013). Effect of diet-induced weight loss on inflammatory cytokines in obese women. J Endocrinol Invest 36, 211–215.

Wang, A., Huen, S.C., Luan, H.H., Yu, S., Zhang, C., Gallezot, J.D., Booth, C.J., and Medzhitov, R. (2016). Opposing Effects of Fasting Metabolism on Tissue Tolerance in Bacterial and Viral Inflammation. Cell 166, 1512–1525 e1512.

Wannemacher, R.W., Jr., Pace, J.G., Beall, R.A., Dinterman, R.E., Petrella, V.J., and Neufeld, H.A. (1979). Role of the liver in regulation of ketone body production during sepsis. The Journal of clinical investigation 64, 1565–1572.

Wei, M., Brandhorst, S., Shelehchi, M., Mirzaei, H., Cheng, C.W., Budniak, J., Groshen, S., Mack, W.J., Guen, E., Di Biase, S., et al. (2017). Fasting-mimicking diet and markers/risk factors for aging, diabetes, cancer, and cardiovascular disease. Science translational medicine 9.

Yamanashi, T., Iwata, M., Kamiya, N., Tsunetomi, K., Kajitani, N., Wada, N., Iitsuka, T., Yamauchi, T., Miura, A., Pu, S., et al. (2017). Beta-hydroxybutyrate, an endogenic NLRP3 inflammasome inhibitor, attenuates stress-induced behavioral and inflammatory responses. Sci Rep 7, 7677.

Youm, Y.H., Nguyen, K.Y., Grant, R.W., Goldberg, E.L., Bodogai, M., Kim, D., D’Agostino, D., Planavsky, N., Lupfer, C., Kanneganti, T.D., et al. (2015). The ketone metabolite beta-hydroxybutyrate blocks NLRP3 inflammasome-mediated inflammatory disease. Nature medicine 21, 263–269.

## References

Amir el, A.D., Davis, K.L., Tadmor, M.D., Simonds, E.F., Levine, J.H., Bendall, S.C., Shenfeld, D.K., Krishnaswamy, S., Nolan, G.P., and Pe’er, D. (2013). viSNE enables visualization of high dimensional single-cell data and reveals phenotypic heterogeneity of leukemia. Nature biotechnology 31, 545– 552.

Brandhorst, S., Wei, M., Hwang, S., Morgan, T.E., and Longo, V.D. (2013). Short-term calorie and protein restriction provide partial protection from chemotoxicity but do not delay glioma progression. Exp Gerontol 48, 1120–1128.

Cignarella, F., Cantoni, C., Ghezzi, L., Salter, A., Dorsett, Y., Chen, L., Phillips, D., Weinstock, G.M., Fontana, L., Cross, A.H., et al. (2018). Intermittent Fasting Confers Protection in CNS Autoimmunity by Altering the Gut Microbiota. Cell metabolism 27, 1222–1235 e1226.

Orr, J.S., Kennedy, A.J., and Hasty, A.H. (2013). Isolation of adipose tissue immune cells. Journal of visualized experiments : JoVE, e50707.

Sergushichev, A.A., Loboda, A.A., Jha, A.K., Vincent, E.E., Driggers, E.M., Jones, R.G., Pearce, E.J., and Artyomov, M.N. (2016). GAM: a web-service for integrated transcriptional and metabolic network analysis. Nucleic acids research 44, W194–200.

Ulland, T.K., Song, W.M., Huang, S.C., Ulrich, J.D., Sergushichev, A., Beatty, W.L., Loboda, A.A., Zhou, Y., Cairns, N.J., Kambal, A., et al. (2017). TREM2 Maintains Microglial Metabolic Fitness in Alzheimer’s Disease. Cell 170, 649–663 e613.

