## Supplemental Figures and Legends for "Dietary intake regulates the circulating inflammatory monocyte pool"

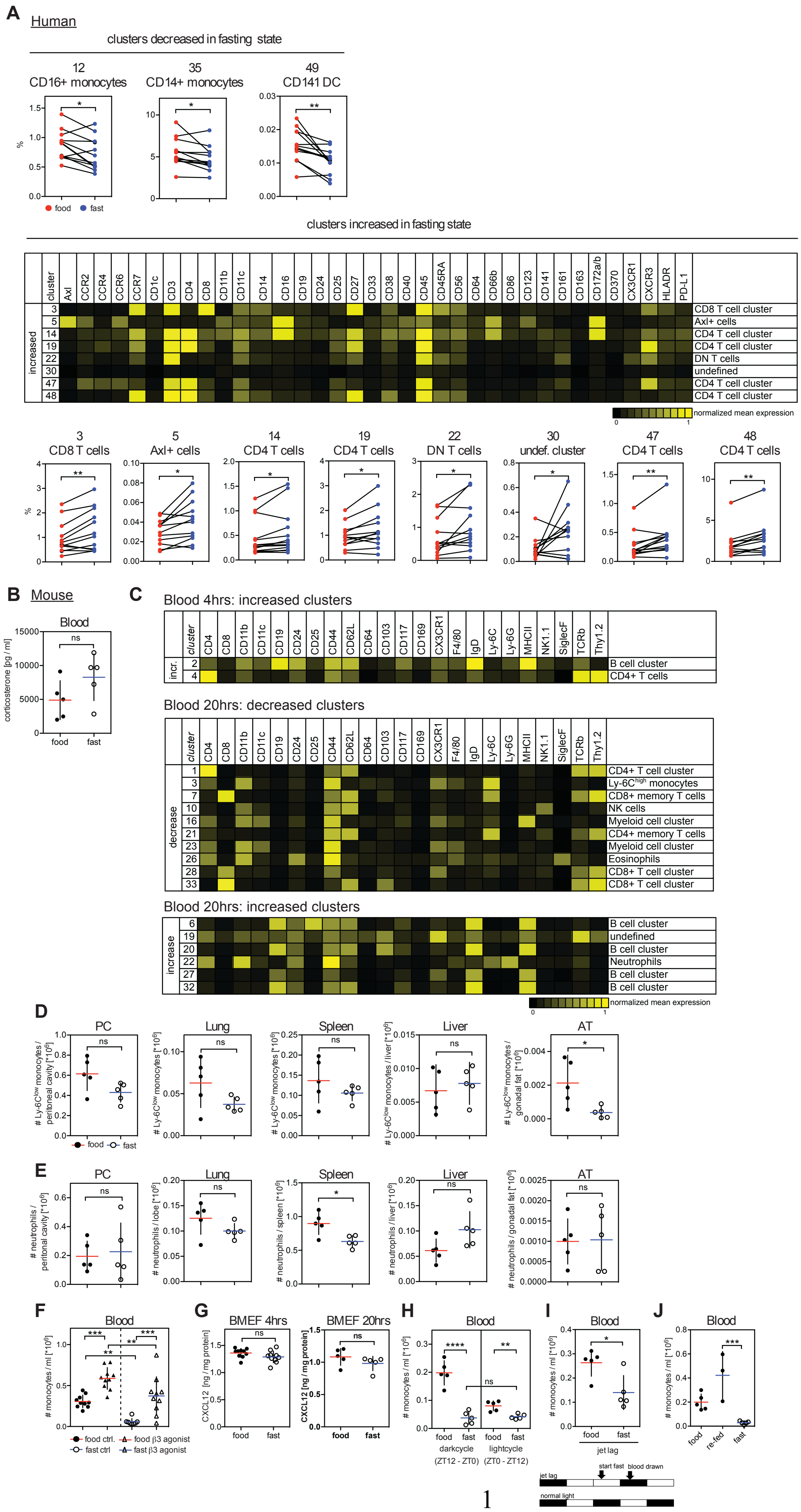

**Figure S1. Profiling of Leukocyte Populations in the Fed and the Fasting State in Healthy Humans and Mice. Related to Figure 1**

(A) Blood was drawn from healthy humans in the fed and in the fasting state and analyzed by CyTOF as described in Figure 1. Multidimensional CyTOF data were clustered using viSNE. Individual representation of each significantly modulated cell cluster. Dots and lines represent paired samples from the fed and fasting state of individual participants. Heatmap shows marker expression on significantly increased clusters. (B) Corticosterone levels in blood of mice that were fed or fasted for 4 hrs. (C) CyTOF analysis of blood from mice that were fed or fasted for 4 hrs and 20 hrs. Heatmaps show mean marker expression on significantly changing cell clusters during fasting. (D and E) Quantification of (D) Ly-6C$^{low}$ monocyte, and (E) neutrophil numbers in peritoneal cavity (PC), lung, spleen, liver, adipose tissue (AT) of mice that were fed or fasted for 20 hrs. (F) Numbers of Ly-6C$^{high}$ monocytes in blood from mice that were fed or fasted, or fed and fasted and gavaged with β3 agonist CL 316,243. Data were pooled from two experiments. (G) Level of CXCL12 in bone marrow extracellular fluid (BMEF) from mice that were fed or fasted for the indicated time. (H) Absolute numbers of Ly-6C$^{high}$ monocytes in the blood of mice that were fasted for 12 hrs either during the light- or the darkcycle. (I) One darkcyle was replaced by a lightcycle prior to analysis as depicted. Absolute numbers of Ly-6C$^{high}$ monocytes in the blood are shown. (J) Absolute numbers of Ly-6C$^{high}$ monocytes in the blood of mice that were fed, fasted overnight or re-fed for 4 hrs after overnight fasting. (B, D to J) Every dot represents one individual animal. Horizontal bar = mean. Vertical bar = SD. Student's t test (A,B,D,E,G to I) or One-way analysis of variance (ANOVA) with Bonferroni's test (F, J) were performed. Statistical significance is indicated by *$P < 0.05$, **$P < 0.01$, ***$P < 0.001$. ns = not significant.

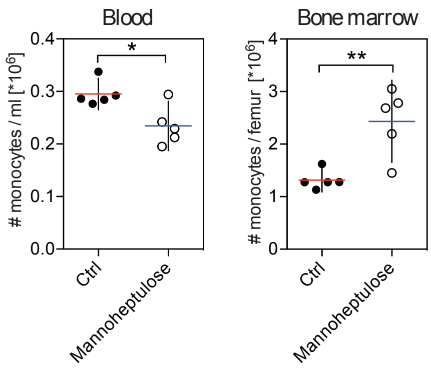


**Figure S2. Inhibition of Glycolysis Reduces Circulating Monocyte Numbers. Related to Figure 2**

Absolute numbers of Ly-6C$^{high}$ monocytes in the blood and bone marrow of mice that were gavaged with water (Ctrl) or D-mannoheptulose once every hour for 4 hrs. Every dot represents one individual animal. Horizontal bar = mean. Vertical bar = SD. Student's t test. Statistical significance is indicated by *P < 0.05, **P < 0.01.

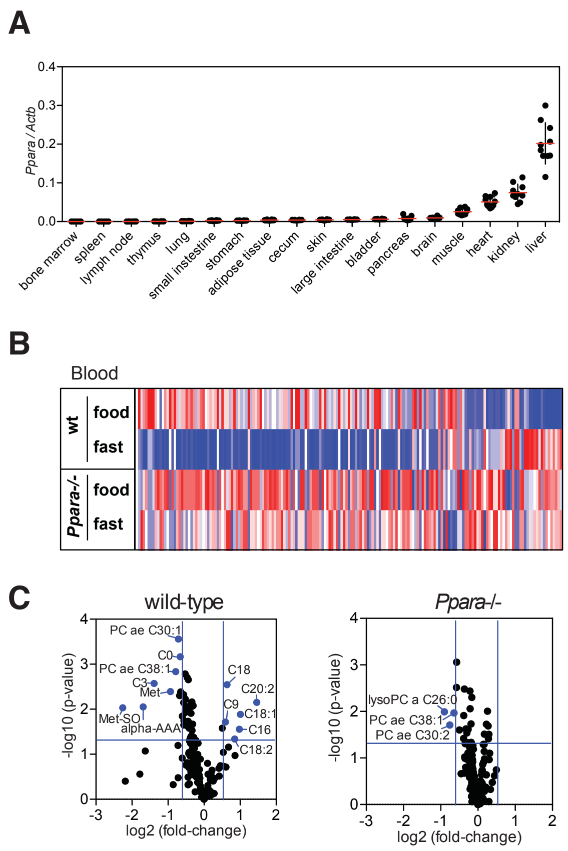


**Figure S3. Profiling of Serum Metabolites in Short-term Fasted Mice. Related to Figure 3**

(A) QPCR for *Ppara* mRNA on indicated tissues. Every dot represents one individual animal. (B and C) 188 metabolites were measured in blood from wt and *Ppara-/-* mice that were fed or fasted for 4 hrs. (B) Heatmap shows z-scores for individual metabolites. (C) Fold-changes and p-values for individual metabolites between fed and fasted wt and *Ppara-/-* mice are represented on a volcano plot.

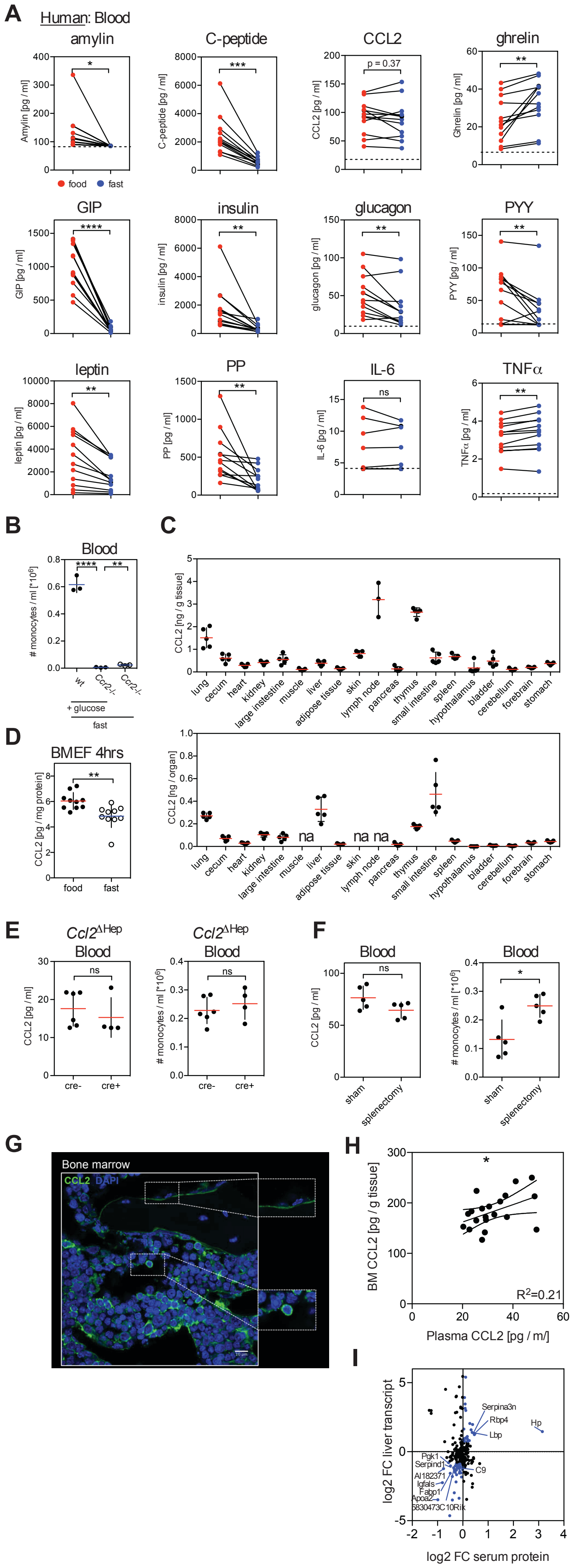


**Figure S4. PPARα Controls Steady-state Blood and Tissue CCL2 Levels. Related to**

**Figure 4**

(A) Multiplex analysis for metabolic hormones in heathy humans. Dots represent paired analysis of individual samples. Dotted lines = limit of detection.

(B) Absolute numbers of Ly-6C$^{high}$ monocytes in fasting wt and *Ccr2*$^{-/-}$ mice that were gavaged with glucose.

(C) Levels of CCL2 protein in indicated tissues.

(D) Level of CCL2 protein in bone marrow extracellular fluid (BMEF).

(E) *Alb*$^{cre/cre}$ mice were crossed to *Ccl2*$^{fl/fl}$ mice to delete *Ccl2* from hepatocytes (*Ccl2*$^{\Delta Hep}$). CCL2 levels and absolute numbers of Ly-6C$^{high}$ monocytes in the blood of cre+ mice deficient in liver CCL2 or cre- littermate controls are shown.

(F) CCL2 levels and absolute numbers of Ly-6C$^{high}$ monocytes in the blood of splenectomized mice.

(G) Representative immunofluorescence of bone marrow sections stained for CCL2 (green) and nuclei (DAPI, blue). Insert zooms in on CCL2 positive cells. Scale bar 10 um.

(H) Plot shows CCL2 production in BM vs. CCL2 levels in plasma.

(I) Significantly changing serum proteins were mapped on differentially expressed genes in hepatocytes upon fasting.

(B to F, H) Every dot represents one individual animal. Horizontal bar = mean. Vertical bar = SD. Student's t test was performed (A, B, D to F). Statistical significance is indicated by *P < 0.05, **P < 0.01, ***P < 0.001, ****P < 0.0001. ns = not significant.

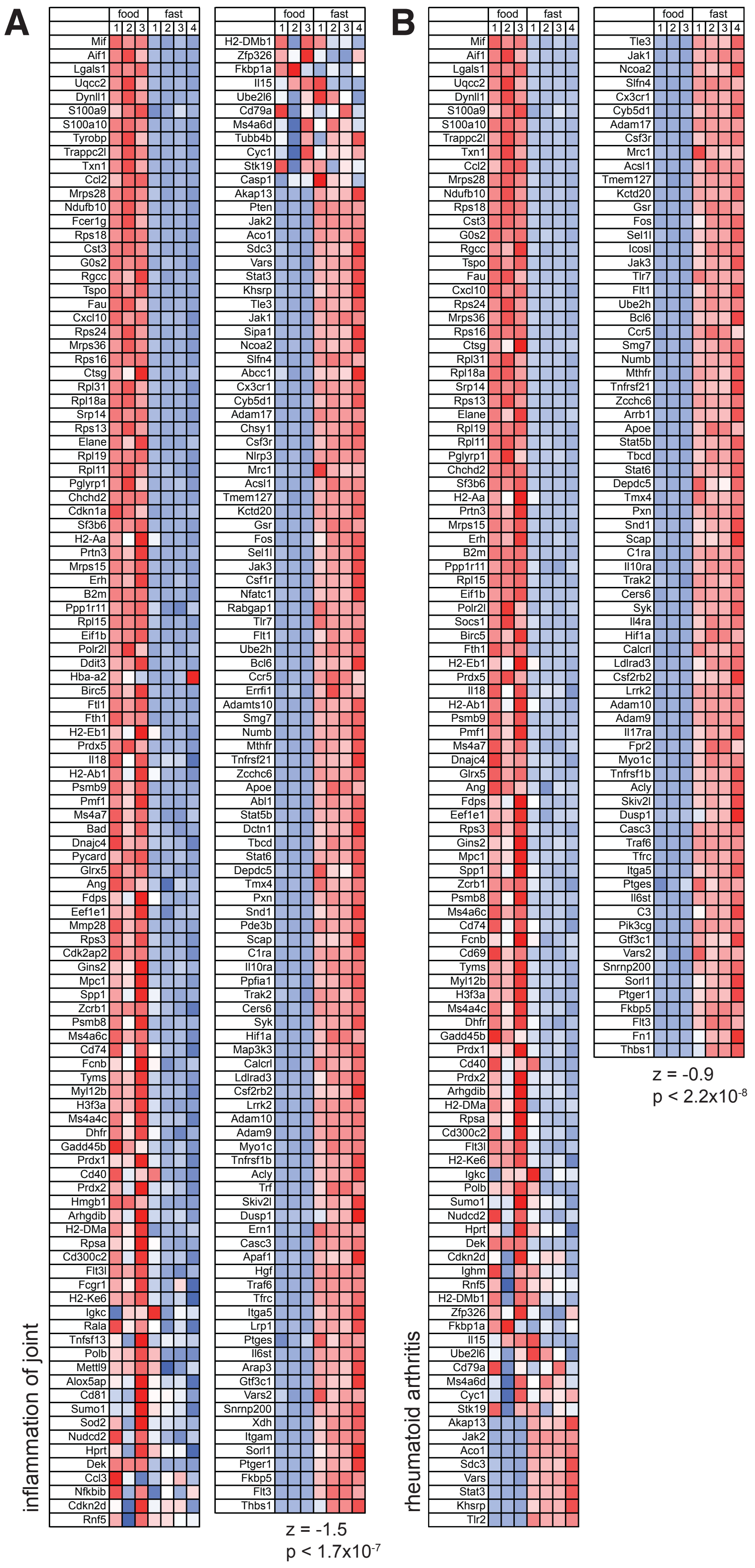


**Figure S5. Fasting Improves Inflammatory Disease Outcome. Related to Figure 6**

(A and B) Differentially expressed genes in monocytes between fed and fasted mice were analyzed using ingenuity pathway analysis (IPA). Heatmaps shows z-scores of genes related to gene sets “inflammation of joint” (A) and “rheumatoid arthritis” (B).
